# THE NEUROPEPTIDE NEUROMEDIN U RECEPTOR NMUR-1 BUFFERS INSULIN RECEPTOR SIGNALING IN BACTERIA-DEPENDENT *C. ELEGANS* SURVIVAL

**DOI:** 10.1101/2025.08.15.670582

**Authors:** Deniz Sifoglu, Bianca Pereira, Christina DeGregory, Rahi Shah, Wolfgang Maier, Joanne Guan, Ian Clark, Dhaval Patel, QueeLim Ch’ng, Joy Alcedo

## Abstract

Distinct microbial environments exert diverse effects on the physiology and survival of the nematode *Caenorhabditis elegans*. Here, we show that *C. elegans* grown on two *Escherichia coli* strains exhibit different survival dynamics. Wild-type *C. elegans* on the B type OP50 exhibit more early deaths compared to *C. elegans* on K-12 type CS180. These early deaths on OP50 are characterized by swollen pharynges (P-deaths) due to bacterial accumulation within the tissue. In contrast, animals on CS180 are more resistant to P-deaths. These bacteria-dependent differences in P-deaths depend on bacterial lipopolysaccharide structures and the activities of the *C. elegans* neuropeptide neuromedin U receptor NMUR-1, which reduces P-deaths on OP50, but not on CS180. Surprisingly, however, NMUR-1 promotes the opposite response when the insulin receptor DAF-2 has reduced function — where NMUR-1 now stimulates P-deaths on OP50, but again with no effect on CS180. We also find that NMUR-1 acts in sensory neurons to promote its bi-directional effects on longevity, which depend on the FOXO transcription factor DAF-16. In addition, NMUR-1 downregulates the expression of the insulin-like peptide *daf-28*, but only when DAF-2 function is not reduced. This suggests a regulatory mechanism through which NMUR-1 maintains insulin receptor DAF-2 signaling at a suitable level. Thus, our studies reveal that NMUR-1 serves to buffer the dynamic range of DAF-2 receptor signaling, thereby optimizing pharyngeal health and survival in response to specific bacteria.

## Introduction

Bacteria are the major dietary source for some animals [1, 2], provide metabolites as part of the intestinal microbiome [3–5], or act as pathogens in other situations [3, 6]. As these three major bacterial functions impact and shape an animal’s life history traits, animals must adapt and respond to their microbial environment for optimal health and survival.

In the nematode worm *C. elegans*, bacteria serve as its primary food source [1], supplying nutrients that stimulate differential gene expression to affect physiology and survival [7–13]. Since *C. elegans* eat different kinds of bacteria, its genetic tractability and the ease of studying its physiology have also made it a useful model for microbiome-derived metabolite studies [5, 14]. At the same time, several bacteria have been shown to pose a threat to *C. elegans* [6]. To isolate the contributions of these bacterial functions, *C. elegans* physiology can be dissected when grown on specific bacterial types — such as wild-type versus mutant bacteria or live versus dead bacteria or different bacterial species [5–13]. These studies reveal that some bacteria are a source of nutrients, metabolites, and/or infection [10, 11, 13, 15].

In the laboratory, *C. elegans* usually feeds on a diet of the B-type *E. coli* OP50 [16], which is not part of the animal’s native microbiome [5]. *E. coli* OP50 has also been shown to be pathogenic to the worm [15, 17]. Colonization of *C. elegans* pharynges by live, proliferating OP50 leads to swelling of the pharynx, ultimately killing the animals — a type of death known as P-death [15, 17]. Presently, the mechanism(s) underlying this type of death remain unclear. For example, the OP50 bacteria-derived cue(s) that promote P-deaths are unknown. Previously, we have shown that worms grown on *E. coli* OP50 live shorter than worms grown on a K-12 type of *E. coli*, CS180, and that this lifespan difference is at least partly dependent on the *E. coli* lipopolysaccharide (LPS) structure [7]. Here we show that LPS structure also mediates OP50-dependent P-deaths in *C. elegans*.

In the host, only a few *C. elegans* genes have been implicated in modulating P-deaths [17–19], which include regulators of innate immunity. Of particular interest is the neuropeptide neuromedin U receptor *nmur-1*, which elicits distinct responses to different pathogenic bacteria [20]. *nmur-1* promotes survival against the pathogen *Enterococcus faecalis*, limits survival on *Salmonella enterica*, and has no effect on *Pseudomonas aeuruginosa* [20]. Interestingly, *nmur-1*, which is expressed in several sensory neurons and in subsets of interneurons and motor neurons [7, 21], has also been shown to mediate the *E. coli* OP50-dependent effects on mitochondrial function and longevity in *C. elegans* [7, 11]. However, the *nmur-1* deletion allele, *ok1387*, used in these studies is also tightly linked to a second mutation, *ot611*, which is located in the gene *filamin-2* (*fln-2*), whose gene product promotes P-deaths [18]. To dissect the effects of *nmur-1* on OP50-dependent P-deaths, we recombined *fln-2(ot611)* away from *nmur-1(ok1387)*.

Here we show that NMUR-1 has complex effects on OP50-dependent *C. elegans* survival but has little or no effect on CS180-dependent worm survival. In the presence of wild-type insulin receptor DAF-2 signaling, the other known regulator of P-deaths [17, 19], wild-type NMUR-1, NMUR-1(+), inhibits P-deaths on OP50. Intriguingly, NMUR-1(+) produces an opposite response on OP50 when DAF-2 insulin receptor function is reduced. In this context, NMUR-1(+) now increases P-deaths, as well as deaths that are not associated with swollen pharynges (non-P deaths). This interaction with DAF-2 suggests that NMUR-1(+), which acts in sensory neurons, adjusts the dynamic range of insulin receptor signaling, a pathway known to be important for survival (reviewed by [22]). We further find that NMUR-1(+) specifically regulates the expression of an insulin-like peptide (ILP) ligand, *daf-28*, when DAF-2 function is unimpaired, which suggests that NMUR-1(+) tunes insulin receptor signaling through defined ILPs. Thus, this mechanism should provide the physiological flexibility necessary in coping with diverse microbial environments.

## Results

### *E. coli* LPS structure modulates *C. elegans* survival dynamics

The *E. coli* B-type OP50 caused worms to live shorter than a K-12 type *E. coli*, CS180 (Fig 1A; Tables 1 and S1; [7]). Worms on OP50 also had a higher rate of early deaths compared to worms on CS180 (Fig 1A), which suggests the presence in OP50 of an early hazard that is absent from CS180. Early deaths on OP50 due to bacterial colonization can be visualized by swollen pharynges (P-deaths; compare Fig 1D to 1F; [15, 17]). Since CS180 reduced deaths in early adulthood (Fig 1A; Tables 1 and S1), we tested if P-deaths contributed to the lifespan differences between wild-type worms on the two bacteria (see Materials and Methods on determination of P-deaths; Fig S1; Table S2). First, we observed that all wild-type P-deaths on OP50 or CS180 occurred by day 15 of adulthood (Fig S2; Table S3). Second, upon censoring all non-P deaths, we found that OP50-fed worms showed about 3 times more P-deaths, when compared to CS180-fed worms (Fig 1B and 1C; Tables 1 and S1), revealing that worms on CS180 were more resistant to P-deaths.

**Fig 1.**
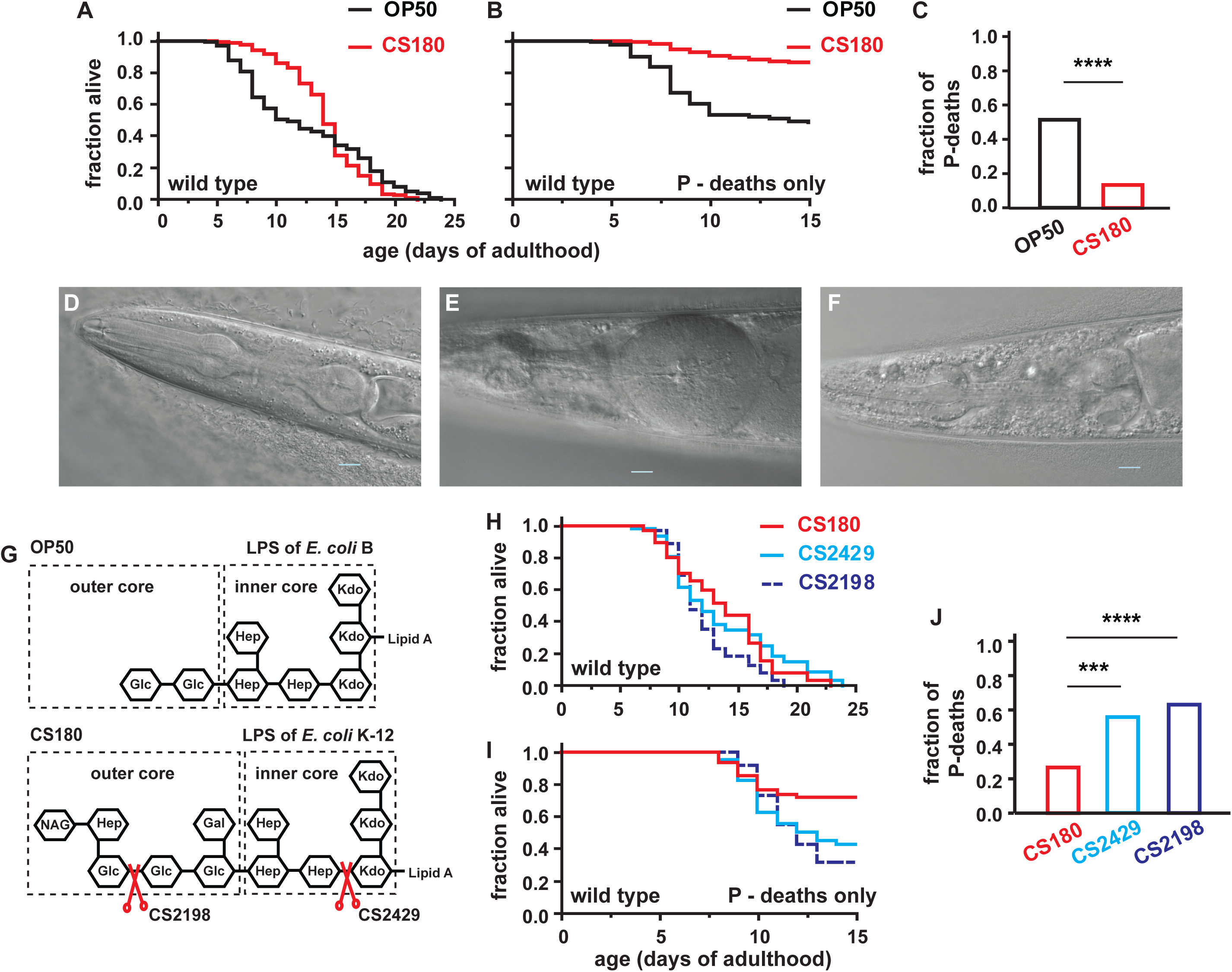
Bacterial food type influences swollen pharynx-dependent deaths in *C. elegans*. **(A)** Wild-type *C. elegans* fed *E. coli* OP50 had more early deaths compared to *C. elegans* fed *E. coli* CS180. **(B)** The early deaths depended on swollen pharynges (P-deaths) in worms fed OP50 compared to worms fed CS180. To generate the survival plots that only depict P-deaths in this figure and subsequent figures, we censored all non-P deaths. **(C)** The fraction of P-deaths out of 144 total deaths on OP50 or 164 total deaths on CS180. **(D)** DIC image of a non-swollen pharynx in a live one-day old adult worm. **(E-F)** DIC images of dead five-day old adult worms that either have a swollen pharynx **(E)** or a non-swollen pharynx **(F)**. Scale bar is 10 μm. **(G)** LPS structures of OP50 and CS180. The red scissors indicate the LPS truncations that correspond to CS2198 and CS2429, which are derived from CS180. **(H-I)** All deaths **(H)** versus P-deaths **(I)** of wild-type *C. elegans* on CS180 and the LPS-truncated CS2198 and CS2429. **(J)** The fraction of P-deaths out of 64 deaths on CS180, 61 deaths on CS2429 and 65 deaths on CS2198. Chi-square analyses were carried out to determine significant differences between the fractions of P-deaths among the different groups of animals on the different bacteria in this figure and subsequent figures. *** denotes *P* < 0.001, whereas **** denotes *P* < 0.0001. See Table 1 for the rest of the statistical analyses that also pertain to this figure and subsequent figures.

**Table 1.** Cumulative statistics of all deaths versus P-deaths on different bacteria. Worms were censored at the time they crawled off the plate, exploded, or bagged, allowing these worms to be incorporated into the data set until the censor date. This avoids loss of information. *P* values that are significant (*P* ≤ 0.05) are italicized and in bold face. If the test population lived longer or had fewer P-deaths than the population to which it is compared, the *P* values are also underlined. Both Wilcoxon and logrank *P* values are shown for comparison (see Materials and Methods on the suitability of one statistical test versus the other). All survival assays were carried out at 25°C on full lawns of the specified bacteria. The superscripts indicate the population to which the test population is compared. The following abbreviations or symbols indicate: WT, wild type; OP, OP50; CS, CS180; *ok1387\**, the genotype *daf-2(e1368)*; *nmur-1(ok1387)*; $, same data used in Figs 2D and 3A; $$, some of the data were also used in Fig 3A; CS24, CS2429; and **, animals carry an extrachromosomal array of *ofm-1p::GFP*.

| Strain/Bacteria:<br>All deaths | Mean<br>Lifespan<br>± SEM<br>(Days) | # Animals<br>Observed/<br>Total Initial<br>Animals<br>(# Trials) | *P* vs specified<br>group<br>(Logrank) | *P* vs specified<br>group<br>(Wilcoxon) | Fig | Strain/Bacteria:<br>P deaths | # Animals<br>Observed/<br>Total Initial<br>Animals<br>(# Trials) | *P* vs specified<br>group<br>(Logrank) | *P* vs specified<br>group<br>(Wilcoxon) | Fig |
| --- | --- | --- | --- | --- | --- | --- | --- | --- | --- | --- |
| *Effects of bacteria on wild-type P deaths* |  |  |  |  |  |  |  |  |  |  |
| Wild type (WT),<br>OP50 | 12.5 ± 0.4 | 142/240<br>(3) | - | - | 1A | Wild type, OP50 | 74/240 (3) | - | - | 1B |
| Wild type (WT),<br>CS180 | 14.2 ± 0.3 | 142/240<br>(3) | 0.34<sup>WT/OP</sup> | <u>< 0.0001<sup>WT/OP</sup></u> | 1A | Wild type,<br>CS180 | 22/240<br>(3) | <u>< 0.0001<sup>WT/OP</sup></u> | <u>< 0.0001<sup>WT/OP</sup></u> | 1B |
| WT, CS180 | 14.0 ± 0.3 | 186/320<br>(3) | - | - |  | WT, CS180 | 26/320<br>(3) | - | - |  |
| WT, CS2429 | 13.9 ± 0.3 | 187/320<br>(3) | 0.94<sup>WT/CS180</sup> | 0.64<sup>WT/CS180</sup> |  | WT, CS2429 | 80/320 (3) | <<br><u>0.0001<sup>WT/CS180</sup></u> | <<br><u>0.0001<sup>WT/CS180</sup></u> |  |
| WT, CS180 | 13.7 ± 0.5 | 64/80 (1) | - | - | 1H | WT, CS180 | 17/80 (1) | - | - | 1I |
| WT, CS2198 | 12.2 ± 0.4 | 65/80 (1) | <u>0.008<sup>WT/CS180</sup></u> | <u>0.05<sup>WT/CS180</sup></u> | 1H | WT, CS2198 | 46/80 (1) | <<br><u>0.0001<sup>WT/CS180</sup></u> | <u>0.004<sup>WT/CS180</sup></u> | 1I |
| WT, CS2429 | 13.7 ± 0.6 | 61/80 (1) | 0.56<sup>WT/CS180</sup> | 0.77<sup>WT/CS180</sup> | 1H | WT, CS2429 | 35/80 (1) | <u>0.004<sup>WT/CS180</sup></u> | <u>0.02<sup>WT/CS180</sup></u> | 1I |
| *nmur-1-dependent P deaths* |  |  |  |  |  |  |  |  |  |  |
| WT, OP50 | 11.9 ± 0.4 | 188/256<br>(1) | - | - | 2A | WT, OP50 | 73/240 (2) | - | - | 2B |
| *nmur-1(ok1387)*,<br>OP50 | 9.8 ± 0.4 | 137/176<br>(1) | <u>0.0005<sup>WT/OP</sup></u> | <u>< 0.0001<sup>WT/OP</sup></u> | 2A | *fln-2(ot611)*,<br>OP50 | 34/230 (2) | <u>< 0.0001<sup>WT/OP</sup></u> | <u>< 0.0001<sup>WT/OP</sup></u> | 2B |
| *fln-2(ot611)*,<br>OP50 | 18.0 ± 0.6 | 49/70 (1) | <u>< 0.0001<sup>WT/OP</sup></u> | <u>< 0.0001<sup>WT/OP</sup></u> | 2A | WT, CS180 | 7/80 (1) | - | - | 2B |
| WT, CS180 | 13.1 ± 0.3 | 117/160<br>(1) | - | - | 2A | *fln-2(ot611)*,<br>CS180 | 0/80 (1) | <u>0.01<sup>WT/CS</sup></u> | <u>0.01<sup>WT/CS</sup></u> | 2B |
| *nmur-1(ok1387)*,<br>CS180 | 14.2 ± 0.4 | 64/80 (1) | <u>0.03<sup>WT/CS</sup></u> | <u>0.02<sup>WT/CS</sup></u> | 2A | WT, OP50 | 177/540<br>(4) | - | - | 2D<sup>\$</sup> |
| *fln-2(ot611)*,<br>CS180 | 12.8 ± 0.3 | 67/80 (1) | 0.34<sup>WT/CS</sup> | 0.50<sup>WT/CS</sup> | 2A | *nmur-1(ok1387)*,<br>OP50 | 244/540<br>(4) | <u>0.0001<sup>WT/OP</sup></u> | <u>< 0.0001<sup>WT/OP</sup></u> | 2D<sup>\$</sup> |
| | | | | | | WT, CS180 | 22/320 (3) | - | - | 2D<sup>\$</sup> |
| | | | | | | *nmur-1(ok1387)*,<br>CS180 | 18/300 (3) | 0.41<sup>WT/CS</sup> | 0.38<sup>WT/CS</sup> | 2D<sup>\$</sup> |
| WT, OP50 | 12.4 ± 0.3 | 210/320<br>(2) | - | - | 2F | WT, OP50 | 108/400<br>(3) | - | - | 2F |
| *nmur-1(lst1672)*,<br>OP50 | 10.4 ± 0.3 | 232/320<br>(2) | <u>< 0.0001<sup>WT/OP</sup></u> | <u>< 0.0001<sup>WT/OP</sup></u> | 2F | *nmur-1(lst1672)*,<br>OP50 | 153/400<br>(3) | <u>< 0.0001<sup>WT/OP</sup></u> | <u>< 0.0001<sup>WT/OP</sup></u> | 2F |

Table 1. Continued
| Strain/Bacteria:<br>All deaths | Mean<br>Lifespan<br>± SEM<br>(Days) | # Animals<br>Observed/<br>Total Initial<br>Animals<br>(# Trials) | *P* vs specified<br>group<br>(Logrank) | *P* vs specified<br>group<br>(Wilcoxon) | Fig | Strain/Bacteria:<br>P deaths | # Animals<br>Observed/<br>Total Initial<br>Animals<br>(# Trials) | *P* vs specified<br>group<br>(Logrank) | *P* vs specified<br>group<br>(Wilcoxon) | Fig |
| --- | --- | --- | --- | --- | --- | --- | --- | --- | --- | --- |
| WT, OP50 | 10.9 ± 0.4 | 117/160<br>(2) | - | - | 3A | WT, OP50 | 177/540<br>(4) | - | - | 3A<sup>\$</sup> |
| *nmur-1(ok1387)*,<br>OP50 | 9.6 ± 0.4 | 120/160<br>(2) | **0.05<sup>WT/OP</sup>** | **0.002<sup>WT/OP</sup>** | 3A | *nmur-1(ok1387)*,<br>OP50 | 244/540<br>(4) | **0.0001<sup>WT/OP</sup>** | **< 0.0001<sup>WT/OP</sup>** | 3A<sup>\$</sup> |
| *daf-2(e1368)*,<br>OP50 | 16.4 ± 0.8 | 94/160 (2) | **< 0.0001<sup>WT/OP</sup>** | **< 0.0001<sup>WT/OP</sup>** | 3A | *daf-2(e1368)*,<br>OP50 | 100/760<br>(4) | **< 0.0001<sup>WT/OP</sup>** | **< 0.0001<sup>WT/OP</sup>** | 3A |
| *daf-2(e1368)*;<br>*nmur-1(ok1387)*,<br>OP50 | 19.1 ± 0.7 | 113/160<br>(2) | **0.03<sup>daf-2/OP</sup>** | **0.001<sup>daf-2/OP</sup>** | 3A | *daf-2(e1368)*;<br>*nmur-1(ok1387)*,<br>OP50 | 56/540 (4) | **< 0.0001<sup>daf-2/OP</sup>** | **< 0.0001<sup>daf-2/OP</sup>** | 3A |
| WT, CS180 | 13.5 ± 0.3 | 106/160<br>(2) | - | - | 3A | WT, CS180 | 22/320 (3) | - | - | 3A<sup>\$</sup> |
| *nmur-1(ok1387)*,<br>CS180 | 14.4 ± 0.3 | 108/140<br>(2) | 0.08<sup>WT/CS</sup> | **0.04<sup>WT/CS</sup>** | 3A | *nmur-1(ok1387)*,<br>CS180 | 18/300 (3) | 0.41<sup>WT/CS</sup> | 0.38<sup>WT/CS</sup> | 3A<sup>\$</sup> |
| *daf-2(e1368)*,<br>CS180 | 23.9 ± 0.5 | 104/160<br>(2) | **< 0.0001<sup>WT/CS</sup>** | **< 0.0001<sup>WT/CS</sup>** | 3A | *daf-2(e1368)*,<br>CS180 | 1/320 (3) | **< 0.0001<sup>WT/CS</sup>** | **< 0.0001<sup>WT/CS</sup>** | 3A |
| *daf-2(e1368)*;<br>*nmur-1(ok1387)*,<br>CS180 | 23.7 ± 0.4 | 105/160<br>(2) | 0.54<sup>daf-2/CS</sup> | 0.42<sup>daf-2/CS</sup> | 3A | *daf-2(e1368)*;<br>*nmur-1(ok1387)*,<br>CS180 | 4/320 (3) | 0.29<sup>daf-2/CS</sup> | 0.29<sup>daf-2/CS</sup> | 3A |
| WT, OP50 | 13.2 ± 0.3 | 145/240<br>(2) | - | - | 3B | WT, OP50 | 63/240 (2) | - | - | 3B |
| *nmur-1(lst1672)*,<br>OP50 | 10.7 ± 0.4 | 153/240<br>(2) | **< 0.0001<sup>WT/OP</sup>** | **< 0.0001<sup>WT/OP</sup>** | 3B | *nmur-1(lst1672)*,<br>OP50 | 89/240 (2) | **< 0.0001<sup>WT/OP</sup>** | **< 0.0001<sup>WT/OP</sup>** | 3B |
| *daf-2(e1368)*,<br>OP50 | 17.8 ± 0.7 | 89/240 (2) | **< 0.0001<sup>WT/OP</sup>** | **< 0.0001<sup>WT/OP</sup>** | 3B | *daf-2(e1368)*,<br>OP50 | 23/240 (2) | **0.003<sup>WT/OP</sup>** | **0.004<sup>WT/OP</sup>** | 3B |
| *daf-2(e1368)*;<br>*nmur-1(lst1672)*,<br>OP50 | 20.5 ± 0.5 | 93/320 (2) | **0.004<sup>daf-2/OP</sup>** | **< 0.0001<sup>daf-2/OP</sup>** | 3B | *daf-2(e1368)*;<br>*nmur-1(lst1672)*,<br>OP50 | 20/320 (2) | **0.009<sup>daf-2/OP</sup>** | **0.002<sup>daf-2/OP</sup>** | 3B |
| WT, OP50 | 10.9 ± 0.4 | 158/220<br>(2) | - | - | 3C | WT, OP50 | 104/220<br>(2) | - | - | 3C |
| *nmur-1(ok1387)*,<br>OP50 | 10.2 ± 0.5 | 140/160<br>(1) | 0.19<sup>WT/OP</sup> | **0.004<sup>WT/OP</sup>** | 3C | *nmur-1(ok1387)*,<br>OP50 | 88/160 (1) | 0.20<sup>WT/OP</sup> | **0.006<sup>WT/OP</sup>** | 3C |
| *daf-2(mu150)*,<br>OP50 | 17.9 ± 0.8 | 130/420<br>(2) | **< 0.0001<sup>WT/OP</sup>** | **< 0.0001<sup>WT/OP</sup>** | 3C | *daf-2(mu150)*,<br>OP50 | 54/420 (2) | **< 0.0001<sup>WT/OP</sup>** | **< 0.0001<sup>WT/OP</sup>** | 3C |
| *daf-2(mu150)*;<br>*nmur-1(ok1387)*,<br>OP50 | 20.7 ± 0.7 | 161/420<br>(2) | **0.004<sup>daf-2/OP</sup>** | **0.0001<sup>daf-2/OP</sup>** | 3C | *daf-2(mu150)*;<br>*nmur-1(ok1387)*,<br>OP50 | 57/420 (2) | 0.06<sup>daf-2/OP</sup> | **0.006<sup>daf-2/OP</sup>** | 3C |

Table 1. Continued
| Strain/Bacteria:<br>All deaths | Mean<br>Lifespan<br>± SEM<br>(Days) | # Animals<br>Observed/<br>Total Initial<br>Animals<br>(# Trials) | *P* vs specified<br>group<br>(Logrank) | *P* vs specified<br>group<br>(Wilcoxon) | Fig | Strain/Bacteria:<br>P deaths | # Animals<br>Observed/<br>Total Initial<br>Animals<br>(# Trials) | *P* vs specified<br>group<br>(Logrank) | *P* vs specified<br>group<br>(Wilcoxon) | Fig |
| --- | --- | --- | --- | --- | --- | --- | --- | --- | --- | --- |
| *Rescue of nmur-1(ok1387) single mutants with nmur-1p::nmur-1* |  |  |  |  |  |  |  |  |  |  |
| WT, OP50 | 11.5 ± 0.5 | 94/160 (1) | - | - | 4A | WT, OP50 | 120/336<br>(2) | - | - | 4A |
| *nmur-1(ok1387)*,<br>OP50 | 10.0 ± 0.4 | 136/160<br>(1) | **0.002<sup>WT/OP</sup>** | **< 0.0001<sup>WT/OP</sup>** | 4A | *nmur-1(ok1387)*,<br>OP50 | 189/336<br>(2) | **0.0002<sup>WT/OP</sup>** | **< 0.0001<sup>WT/OP</sup>** | 4A |
| *nmur-1(ok1387)*;<br>*nmur-1p::nmur-1*,<br>OP50 | 12.3 ± 0.4 | 105/160<br>(1) | 0.22<sup>WT/OP</sup><br>**< 0.0001<sup>ok1387</sup>** | **0.05<sup>WT/OP</sup>**<br>**< 0.0001<sup>ok1387</sup>** | 4A | *nmur-1(ok1387)*;<br>*nmur-1p::nmur-1*,<br>OP50 | 120/336<br>(2) | **0.02<sup>WT/OP</sup>**<br>**< 0.0001<sup>ok1387</sup>** | **0.04<sup>WT/OP</sup>**<br>**< 0.0001<sup>ok1387</sup>** | 4A |
| *Rescue of nmur-1(ok1387) single mutants with osm-6p::nmur-1* |  |  |  |  |  |  |  |  |  |  |
| WT, OP50 | 11.0 ± 0.4 | 91/176 (1) | - | - | 4B | WT, OP50 | 129/336<br>(2) | - | - | 4B |
| *nmur-1(ok1387)*,<br>OP50 | 9.5 ± 0.3 | 118/165<br>(1) | **0.0008<sup>WT/OP</sup>** | **< 0.0001<sup>WT/OP</sup>** | 4B | *nmur-1(ok1387)*,<br>OP50 | 178/325<br>(2) | **< 0.0001<sup>WT/OP</sup>** | **< 0.0001<sup>WT/OP</sup>** | 4B |
| *nmur-1(ok1387)*;<br>*osm-6p::nmur-1*,<br>OP50 | 10.5 ± 0.4 | 90/176 (1) | 0.41<sup>WT/OP</sup><br>**0.01<sup>ok1387</sup>** | 0.35<sup>WT/OP</sup><br>**0.0002<sup>ok1387</sup>** | 4B | *nmur-1(ok1387)*;<br>*osm-6p::nmur-1*,<br>OP50 | 89/336 (2) | **0.006<sup>WT/OP</sup>**<br>**< 0.0001<sup>ok1387</sup>** | **0.005<sup>WT/OP</sup>**<br>**< 0.0001<sup>ok1387</sup>** | 4B |
| *Rescue of daf-2(e1368); nmur-1(ok1387) double mutants with nmur-1p::nmur-1* |  |  |  |  |  |  |  |  |  |  |
| WT, OP50 | 11.3 ± 0.5 | 92/160 (1) | - | - | 4C | WT, OP50 | 120/416<br>(3) | - | - | 4C |
| *daf-2(e1368)*,<br>OP50 | 17.2 ± 1.0 | 71/160 (1) | **< 0.0001<sup>WT/OP</sup>** | **< 0.0001<sup>WT/OP</sup>** | 4C | *daf-2(e1368)*,<br>OP50 | 55/416 (3) | **< 0.0001<sup>WT/OP</sup>** | **< 0.0001<sup>WT/OP</sup>** | 4C |
| *daf-2(e1368)*;<br>*nmur-1(ok1387)*,<br>OP50 | 19.8 ± 0.8 | 75/160 (1) | 0.15<sup>daf-2/OP</sup> | **0.005<sup>daf-2/OP</sup>** | 4C | *daf-2(e1368)*;<br>*nmur-1(ok1387)*,<br>OP50 | 55/496 (3) | **0.007<sup>daf-2/OP</sup>** | **0.0002<sup>daf-2/OP</sup>** | 4C |
| *daf-2(e1368)*;<br>*nmur-1(ok1387)*;<br>*nmur-1p::nmur-1*,<br>OP50 | 17.0 ± 0.8 | 74/160 (1) | 0.45<sup>daf-2/OP</sup><br>**0.01<sup>ok1387\*</sup>** | 0.95<sup>daf-2/OP</sup><br>**0.002<sup>ok1387\*</sup>** | 4C | *daf-2(e1368)*;<br>*nmur-1(ok1387)*;<br>*nmur-1p::nmur-1*,<br>OP50 | 50/496 (3) | 0.06<sup>daf-2/OP</sup><br>0.6<sup>ok1387*</sup> | 0.22<sup>daf-2/OP</sup><br>**0.05<sup>ok1387\*</sup>** | 4C |
| *Rescue of daf-2(e1368); nmur-1(ok1387) double mutants with osm-6p::nmur-1* |  |  |  |  |  |  |  |  |  |  |
| WT, OP50 | 11.2 ± 0.3 | 183/336<br>(2) | - | - | 4D | WT, OP50 | 98/336 (2) | - | - | 4D |
| *daf-2(e1368)*,<br>OP50 | 16.9 ± 0.6 | 150/336<br>(2) | **< 0.0001<sup>WT/OP</sup>** | **< 0.0001<sup>WT/OP</sup>** | 4D | *daf-2(e1368)*,<br>OP50 | 41/336 (2) |  |  | 4D |
| *daf-2(e1368)*;<br>*nmur-1(ok1387)*,<br>OP50 | 18.8 ± 0.5 | 164/336<br>(2) | **0.04<sup>daf-2/OP</sup>** | **0.0003<sup>daf-2/OP</sup>** | 4D | *daf-2(e1368)*;<br>*nmur-1(ok1387)*,<br>OP50 | 33/336 (2) | **0.05<sup>daf-2/OP</sup>** | **0.01<sup>daf-2/OP</sup>** | 4D |

Table 1. Continued
| Strain/Bacteria:<br>All deaths | Mean<br>Lifespan<br>± SEM<br>(Days) | # Animals<br>Observed/<br>Total Initial<br>Animals<br>(# Trials) | *P* vs specified<br>group<br>(Logrank) | *P* vs specified<br>group<br>(Wilcoxon) | Fig | Strain/Bacteria:<br>P deaths | # Animals<br>Observed/<br>Total Initial<br>Animals<br>(# Trials) | *P* vs specified<br>group<br>(Logrank) | *P* vs specified<br>group<br>(Wilcoxon) | Fig |
| --- | --- | --- | --- | --- | --- | --- | --- | --- | --- | --- |
| *daf-2(e1368);*<br>*nmur-1(ok1387);*<br>*osm-6p::nmur-1,*<br>*OP50* | 15.8 ± 0.7 | 130/336<br>(2) | 0.33<sup>daf-2/OP</sup><br>**0.006**<sup>ok1387*</sup> | **0.05**<sup>daf-2/OP</sup><br>**< 0.0001**<sup>ok1387*</sup> | 4D | *daf-2(e1368);*<br>*nmur-1(ok1387);*<br>*osm-6p::nmur-1,*<br>*OP50* | 21/336 (2) | 0.16<sup>daf-2/OP</sup><br>0.65<sup>ok1387*</sup> | 0.08<sup>daf-2/OP</sup><br>0.48<sup>ok1387*</sup> | 4D |
| *LPS-dependence* |  |  |  |  |  |  |  |  |  |  |
| WT, CS180 | 13.9 ± 0.3 | 172/320<br>(3) | - | - | 5A<sup>\$\$</sup> | WT, CS180 | 22/320 (3) | - | - | 5A<sup>\$\$</sup> |
| *nmur-1(ok1387),*<br>*CS180* | 14.7 ± 0.3 | 185/300<br>(3) | 0.06<sup>WT/CS</sup> | **0.02**<sup>WT/CS</sup> | 5A<sup>\$\$</sup> | *nmur-1(ok1387),*<br>*CS180* | 18/300 (3) | 0.41<sup>WT/CS</sup> | 0.38<sup>WT/CS</sup> | 5A<sup>\$\$</sup> |
| *daf-2(e1368),*<br>*CS180* | 23.9 ± 0.5 | 104/160<br>(2) | **< 0.0001**<sup>WT/CS</sup> | **< 0.0001**<sup>WT/CS</sup> | 5A<sup>\$\$</sup> | *daf-2(e1368),*<br>*CS180* | 1/320 (3) | **< 0.0001**<sup>WT/CS</sup> | **< 0.0001**<sup>WT/CS</sup> | 5A<sup>\$\$</sup> |
| *daf-2(e1368);*<br>*nmur-1(ok1387),*<br>*CS180* | 23.7 ± 0.4 | 105/160<br>(2) | 0.54<sup>daf-2/CS</sup> | 0.54<sup>daf-2/CS</sup> | 5A<sup>\$\$</sup> | *daf-2(e1368);*<br>*nmur-1(ok1387),*<br>*CS180* | 4/320 (3) | 0.29<sup>daf-2/CS</sup> | 0.29<sup>daf-2/CS</sup> | 5A<sup>\$\$</sup> |
| WT, CS2429 | 14.6 ± 0.3 | 126/240<br>(2) | - | - | 5B | WT, CS2429 | 87/400 (3) | - | - | 5B |
| *nmur-1(ok1387),*<br>*CS2429* | 12.9 ± 0.3 | 158/220<br>(2) | **0.005**<sup>WT/CS24</sup> | **0.0004**<sup>WT/CS24</sup> | 5B | *nmur-1(ok1387),*<br>*CS2429* | 127/380<br>(3) | **0.0002**<sup>WT/CS24</sup> | **< 0.0001**<sup>WT/CS24</sup> | 5B |
| *daf-2(e1368),*<br>*CS2429* | 25.4 ± 0.7 | 58/82 (1) | **< 0.0001**<sup>WT/CS24</sup> | **< 0.0001**<sup>WT/CS24</sup> | 5B | *daf-2(e1368),*<br>*CS2429* | 33/400 (3) | **< 0.0001**<sup>WT/CS24</sup> | **< 0.0001**<sup>WT/CS24</sup> | 5B |
| *daf-2(e1368);*<br>*nmur-1(ok1387),*<br>*CS2429* | 26.7 ± 0.6 | 50/80 (1) | 0.55<sup>daf-2/CS24</sup> | 0.27<sup>daf-2/CS24</sup> | 5B | *daf-2(e1368);*<br>*nmur-1(ok1387),*<br>*CS2429* | 16/400 (3) | **0.005**<sup>daf-2/CS24</sup> | **0.004**<sup>daf-2/CS24</sup> | 5B |
| *daf-28 survival phenotypes\*\** |  |  |  |  |  |  |  |  |  |  |
| WT, OP50 | 11.3 ± 0.3 | 174/211<br>(2) | - | - |  | WT, OP50 | 99/211 (2) | - | - | 6D |
| *daf-28(tm2308),*<br>*OP50* | 11.6 ± 0.4 | 158/223<br>(2) | 0.41<sup>Ctrl/OP</sup> | 0.48<sup>WT/OP</sup> |  | *daf-28(tm2308),*<br>*OP50* | 72/223 (2) | 0.06<sup>WT/OP</sup> | **0.05**<sup>Ctrl/OP</sup> | 6D |
| *daf-16-dependence* |  |  |  |  |  |  |  |  |  |  |
| WT, OP50 | 11.6 ± 0.5 | 136/176<br>(1) | - | - | 7A,B | WT, OP50 | 138/352<br>(2) | - | - | 7A,B |
| *nmur-1(ok1387),*<br>*OP50* | 9.8 ± 0.4 | 138/176<br>(1) | **0.003**<sup>WT/OP</sup> | **< 0.0001**<sup>WT/OP</sup> | 7A | *nmur-1(ok1387),*<br>*OP50* | 149/352<br>(2) | **0.02**<sup>WT/OP</sup> | **0.0002**<sup>WT/OP</sup> | 7A |
| *daf-16(mu86),*<br>*OP50* | 9.1 ± 0.3 | 143/176<br>(1) | **< 0.0001**<sup>WT/OP</sup> | **0.0001**<sup>WT/OP</sup> | 7A | *daf-16(mu86),*<br>*OP50* | 140/352<br>(2) | 0.08<sup>WT/OP</sup> | **0.0007**<sup>WT/OP</sup> | 7A |

Table 1. Continued
| Strain/Bacteria:<br>All deaths | Mean<br>Lifespan<br>± SEM<br>(Days) | # Animals<br>Observed/<br>Total Initial<br>Animals<br>(# Trials) | *P* vs specified<br>group<br>(Logrank) | *P* vs specified<br>group<br>(Wilcoxon) | Fig | Strain/Bacteria:<br>P deaths | # Animals<br>Observed/<br>Total Initial<br>Animals<br>(# Trials) | *P* vs specified<br>group<br>(Logrank) | *P* vs specified<br>group<br>(Wilcoxon) | Fig |
| --- | --- | --- | --- | --- | --- | --- | --- | --- | --- | --- |
| *daf-16(mu86);*<br>*nmur-1(ok1387),*<br>*OP50* | 9.2 ± 0.3 | 142/176<br>(1) | 0.50<sup>*daf-16/OP*</sup> | 0.83<sup>*daf-16/OP*</sup> | 7A,B | *daf-16(mu86);*<br>*nmur-1(ok1387),*<br>*OP50* | 149/352<br>(2) | 0.89<sup>*daf-16/OP*</sup> | 0.75<sup>*daf-16/OP*</sup> | 7A,B |
| *daf-2(e1368),*<br>*OP50* | 18.4 ± 0.7 | 109/176<br>(1) | **< 0.0001<sup>WT/OP</sup>** | **< 0.0001<sup>WT/OP</sup>** | 7B | *daf-2(e1368),*<br>*OP50* | 58/352 (2) | **< 0.0001<sup>WT/OP</sup>** | **< 0.0001<sup>WT/OP</sup>** | 7B |
| *daf-2(e1368);*<br>*nmur-1(ok1387),*<br>*OP50* | 21.6 ± 0.6 | 116/176<br>(1) | **0.008<sup>*daf-2/OP*</sup>** | **0.0004<sup>*daf-2/OP*</sup>** | 7B | *daf-2(e1368);*<br>*nmur-1(ok1387),*<br>*OP50* | 43/352 (2) | **0.02<sup>*daf-2/OP*</sup>** | **0.003<sup>*daf-2/OP*</sup>** | 7B |
| *daf-16(mu86);*<br>*daf-2(e1368),*<br>*OP50* | 8.9 ± 0.3 | 120/176<br>(1) | 0.70<sup>*daf-16/OP*</sup> | 0.44<sup>*daf-16/OP*</sup> | 7B | *daf-16(mu86);*<br>*daf-2(e1368),*<br>*OP50* | 146/352<br>(2) | 0.22<sup>*daf-16/OP*</sup> | 0.27<sup>*daf-16/OP*</sup> | 7B |
| *daf-16(mu86);*<br>*daf-2(e1368);*<br>*nmur-1(ok1387),*<br>*OP50* | 9.0 ± 0.3 | 140/176<br>(1) | 0.78<sup>*daf-16/OP*</sup> | 0.88<sup>*daf-16/OP*</sup> | 7B | *daf-16(mu86);*<br>*daf-2(e1368);*<br>*nmur-1(ok1387),*<br>*OP50* | 161/352<br>(2) | 0.19<sup>*daf-16/OP*</sup> | 0.35<sup>*daf-16/OP*</sup> | 7B |

Next, we asked what bacterial cues might contribute to the P-death differences between OP50-grown and CS180-grown worms. We previously showed that the *E. coli* LPS structure can modulate *C. elegans* longevity [7]. CS180 LPS truncation mutants, CS2198 and CS2429 (Fig 1G), have been shown to shorten wild-type worm lifespan [7]. Hence, we compared the number of P-deaths on both CS2198 and CS2429 to those on CS180. While we found that only the short LPS mutant *E. coli* CS2198 decreased worm lifespan in this study (Fig 1H; Tables 1 and S1), both *E. coli* strains with the shorter LPS produced more P-deaths than CS180 (Fig 1I and 1J; Tables 1 and S1). Thus, our results show that altering the core LPS structure is sufficient to promote *E. coli* colonization of the pharynx and increased P-deaths.

### Opposing effects of *nmur-1* on *E. coli* OP50 depends on the *daf-2* insulin receptor

We then asked what host genetic factors influence the bacterial-dependent P-deaths. One candidate gene is the neuropeptide neuromedin U receptor *nmur-1*, which has been demonstrated to mediate bacteria-specific innate immune responses [20], some of which might depend on the LPS structure of *E. coli* [7]. The *nmur-1(ok1387)* deletion mutation used in these studies is tightly linked to the *ot611* mutation present in the putative actin-binding scaffold protein gene *fln-2*, which also regulates P-deaths [18]. To address OP50-dependent P-death phenotypes that are specific to the *nmur-1* deletion, we separated the *fln-2(ot611)* and *nmur-1(ok1387)* mutations.

This approach enabled us to dissect the complex effects of the isolated *nmur-1(ok1387)* mutation. First, *nmur-1(ok1387)* produced a short lifespan on OP50 but not on CS180, whereas the *fln-2(ot611)* mutant lived long only on OP50 (Fig 2A; Tables 1 and S4). While *fln-2(ot611)* mutants also had fewer P-deaths (Fig 2B and 2C; Tables 1 and S4), the *nmur-1(ok1387)* mutant had more P-deaths on OP50 but not on CS180 (Fig 2D and 2E; Tables 1 and S4). These results were recapitulated in a second independent deletion allele of *nmur-1*, *lst1672* ([21]; Fig 2F and 2G; Tables 1 and S4). Loss of *nmur-1* affected survival largely through pharynx-dependent deaths (Fig S3A to S3B; Tables S5 and S6). When we only counted deaths that are characterized by unswollen pharynges (non-P deaths; Fig S3C; Tables S5 and S6), this time censoring all P-deaths, the survival of *nmur-1* mutants is more similar to wild-type survival. In this context, *nmur-1(+)* acts to protect *C. elegans* from P-deaths in a bacterial-dependent manner.

**Fig 2.**
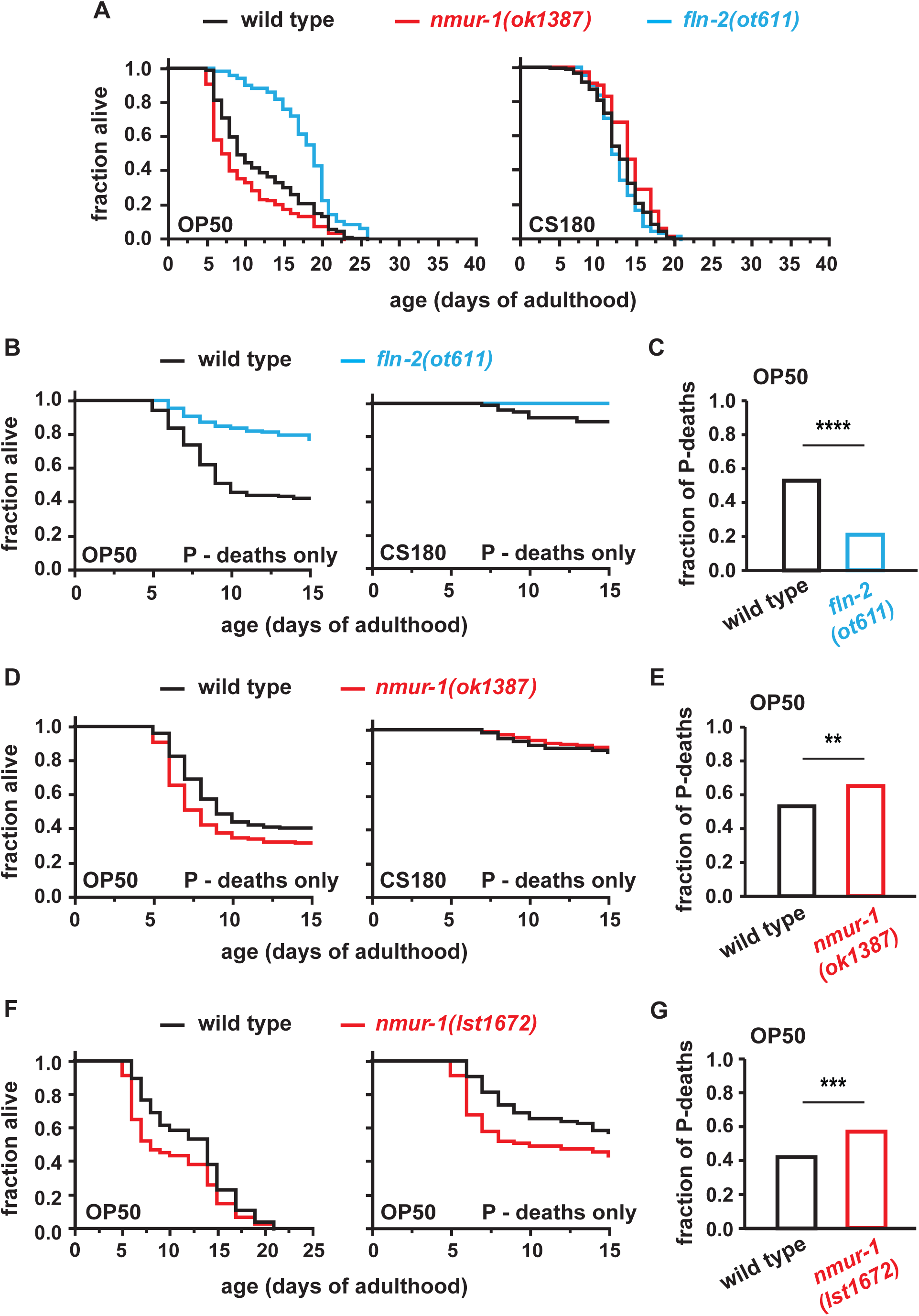
Wild-type *nmur-1* decreases pharynx-dependent deaths in a bacteria-dependent manner. **(A)** On OP50, the *nmur-1*(*ok1387*) single mutant shortened lifespan whereas the *fln-2(ot611)* single mutant extended lifespan. On CS180, *nmur-1* mutants lived slightly longer than wild type, but *fln-2* mutants lived like wild type. **(B)** *fln-2(ot611)* had fewer P-deaths on OP50. **(C)** The fraction of P-deaths on OP50 out of 138 deaths for wild type and 162 deaths for *fln-2(ot611)*. **(D)** *nmur-1*(*ok1387*) had more P-deaths on OP50 but not on CS180. **(E)** The fraction of P-deaths on OP50 out of 203 deaths for wild type and 231 deaths for *nmur-1(ok1387)*. **(F)** A second allele of *nmur-1*, *lst1672*, also shortened lifespan and increased P deaths on OP50. **(G)** The fraction of P-deaths on OP50 out of 257 deaths for wild type and 269 deaths for *nmur-1(lst1672)*. The following symbols denote: **, *P* < 0.01; ***, *P* < 0.001; and ****, *P* < 0.0001.

Intriguingly, the effect of *nmur-1* deletion on OP50-dependent deaths is altered by the presence of mutations that reduce DAF-2 protein activity. The wild-type insulin receptor DAF-2 promotes deaths caused by bacterial colonization and pharyngeal swelling [17, 19]. In insects and mammals, neuromedin U signaling influences insulin signaling by suppressing insulin secretion under certain contexts [23–27], which led us to test whether the P-death phenotype of *nmur-1* mutations would be *daf-2*-dependent. Unexpectedly, the *daf-2(e1368)* mutation, which decreases receptor protein function, not only lengthened lifespan and suppressed P-deaths but also revealed that the *nmur-1* mutations have bi-directional effects on lifespan and P-deaths. Unlike animals with wild-type DAF-2 function (Fig. 2A and 2D to 2G; Tables 1 and S4), *nmur-1(ok1387)* and *nmur-1(lst1672)* now led to fewer P-deaths in *daf-2* reduction-of-function mutant backgrounds (*e1368* or *mu150*) on OP50 (Fig 3; Tables 1 and S7), but not on CS180 (Fig 3A; Tables 1 and S7). Thus, deletion of *nmur-1* further extends the long lifespan of *daf-2* mutants in a bacteria-dependent manner (Fig 3; Tables 1 and S7).

**Fig 3.**
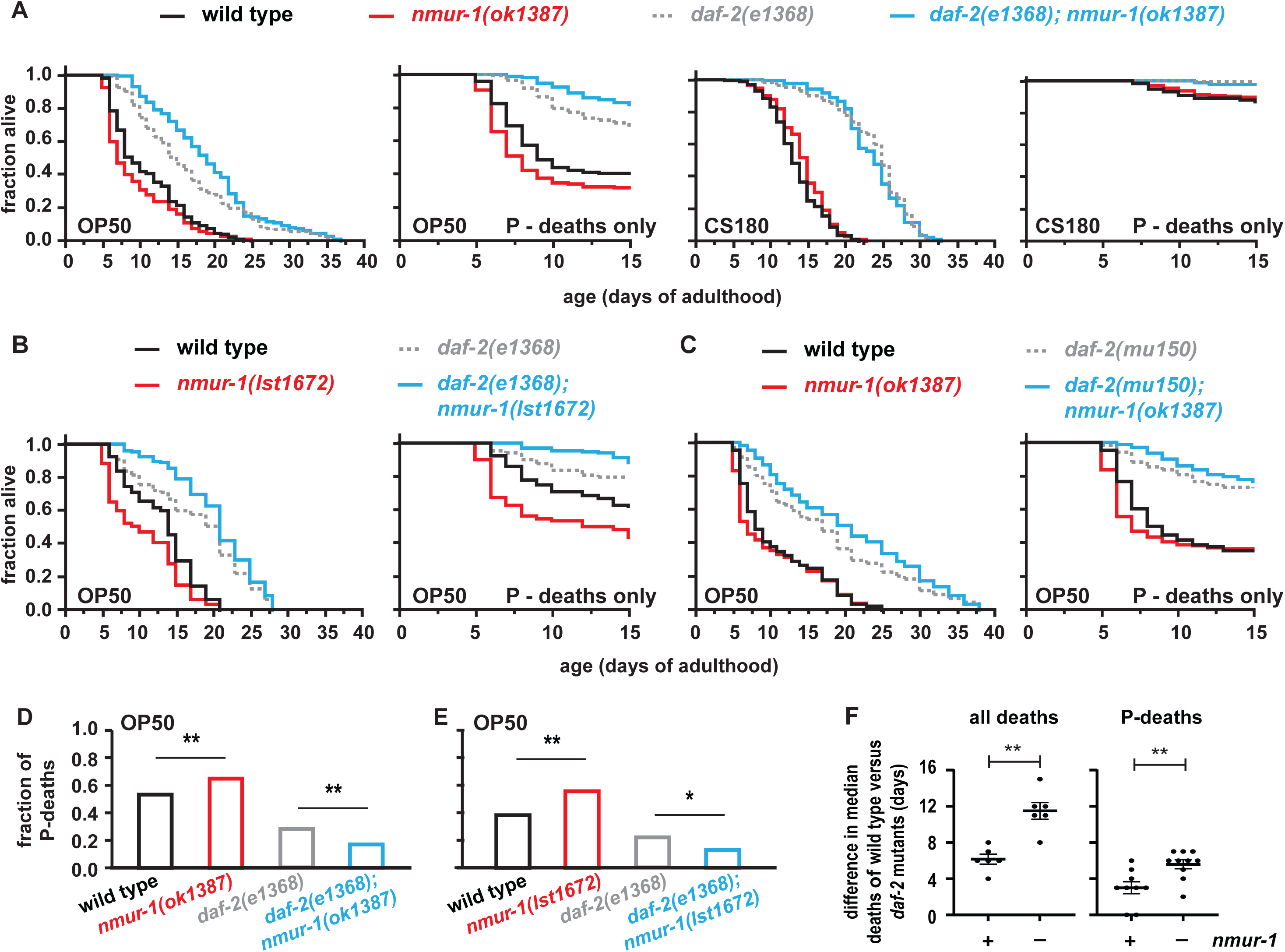
In contrast, under lowered DAF-2 insulin receptor activity, wild-type *nmur-1* increases P-deaths. **(A)** In the *daf-2(e1368)* reduction-of-function mutant background, *nmur-1(ok1387)* had an opposing effect on lifespan on OP50—*nmur-1(ok1387)* increased longevity by decreasing P-deaths. Again, *nmur-1(ok1387)* had little effect on CS180. **(B)** A second allele of *nmur-1*, *lst1672*, also increased lifespan and reduced P-deaths in the *daf-2(e1368)* mutant background. **(C)** *nmur-1(ok1387)* also enhanced the lifespan of another reduction-of-function allele of *daf-2*, *mu150*, which led to fewer P-deaths. **(D)** The fraction of P-deaths on OP50 out of 203 deaths for wild type, 231 deaths for *nmur-1(ok1387)* single mutants, 182 deaths for *daf-2(e1368)* single mutants and 206 deaths for *daf-2(e1368)*; *nmur-1(ok1387)* double mutants out of 2 trials. **(E)** The fraction of P-deaths on OP50 out of 165 deaths for wild type, 160 deaths for *nmur-1(lst1672)* single mutants, 105 deaths for *daf-2(e1368)* single mutants and 156 deaths for *daf-2(e1368)*; *nmur-1(lst1672)* double mutants out of 2 trials. **(F)** *Left panel*: The difference in the median time of all deaths between wild type and *daf-2* reduction-of-function mutants in the presence (n = 6 trials) or absence of *nmur-1* (n = 6 trials). *Right panel*: The difference in the median time of P-deaths between wild type and *daf-2* reduction-of-function mutants in the presence (n = 9 trials) or absence of *nmur-1* (n = 10 trials). Significance in median differences is determined by the Mann-Whitney test. The following symbols denote: *, *P* < 0.05; **, *P* < 0.01.

In contrast to animals with wild-type *daf-2*, the *nmur-1* mutation modulated both P-deaths and non-P deaths in a *daf-2* reduction-of-function mutant background (Figs S3D to S3F, S4A and S4C; Tables S5, S6 and S8). This suggests that *nmur-1(+)* affects survival through other mechanisms besides pharyngeal colonization. Importantly, the opposing *nmur-1* mutant phenotypes in the wild-type *daf-2* versus mutant *daf-2* backgrounds suggest that NMUR-1(+) adjusts and buffers insulin receptor protein activity. Specifically, loss of *nmur-1* enhances the impact of *daf-*2 mutations on lifespan: it increases the difference in the median time of death between wild type and *daf-2* reduction-of-function mutants, when considering either total deaths or P-deaths (Fig 3F; Table S7). This increase in the dynamic range of median lifespan between wild-type and *daf-2* mutants implies that loss of *nmur-1* leads to an animal’s greater sensitivity to DAF-2 protein activity levels. Together, these results suggest a role for wild-type NMUR-1 in buffering the impact of DAF-2 receptor activity on lifespan.

### *nmur-1* promotes opposing effects on survival by acting in sensory neurons in response to LPS structure

To address the mechanisms through which *nmur-1* exerts its multiple activities, we first verified its role through rescue experiments. Expression of *nmur-1(+)* from its own promoter [21] rescued the *nmur-1* short-lived single mutant phenotype (Fig 4A; Tables 1 and S9). When *daf-2* activity is reduced, extrachromosomal expression of *nmur-1(+)* from its own promoter also rescued the longer life phenotype due to the *nmur-1* mutation (Fig 4C; Tables 1 and S9). The same construct rescued the P-death (Fig 4C; Tables 1 and S9) and non-P death phenotypes caused by *nmur-1(ok1387)* in the *daf-2(e1368)* background (Fig S4A and S4B; Tables S5 and S8).

**Fig 4.**
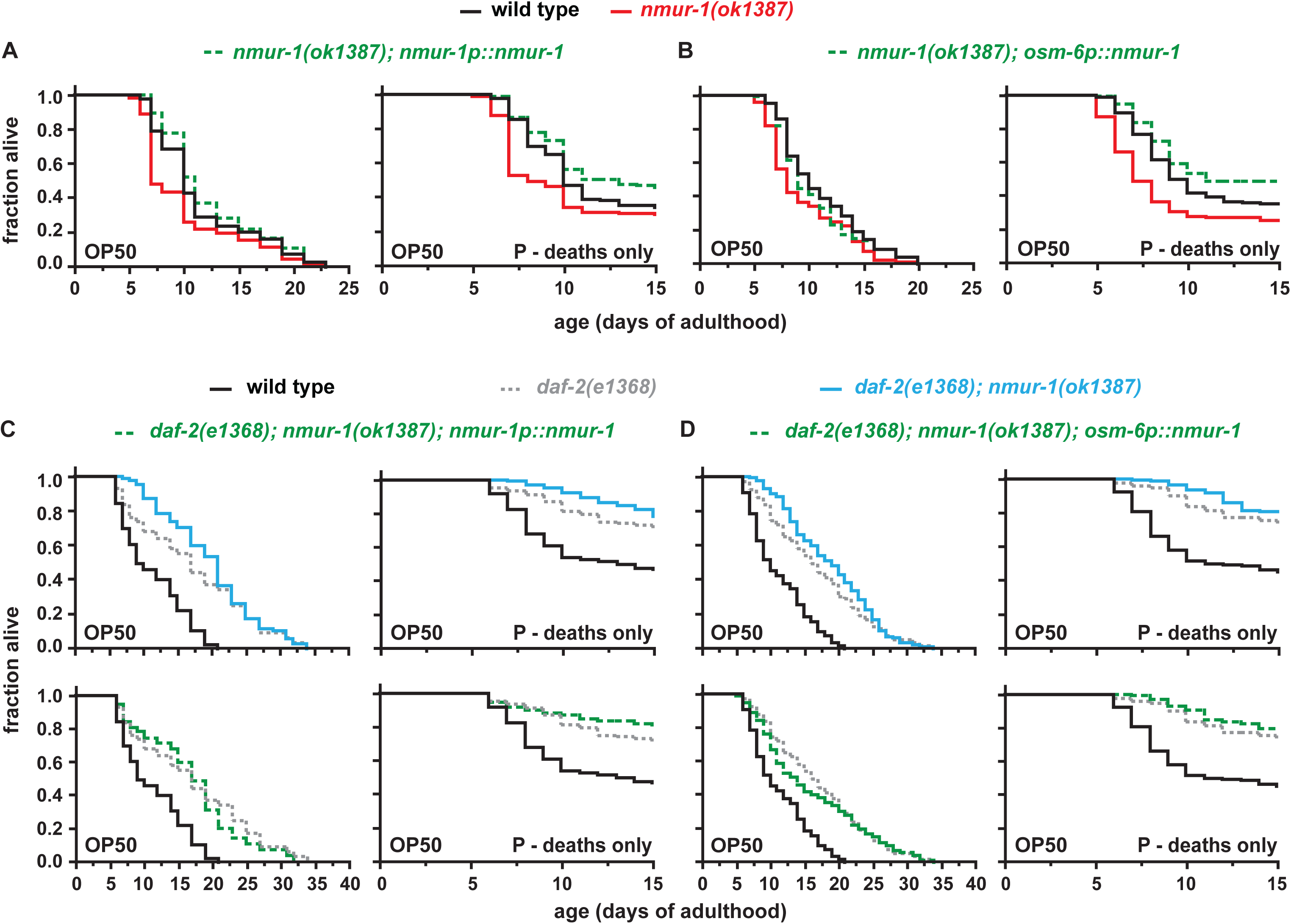
*nmur-1* acts in sensory neurons to modulate longevity and P deaths. **(A-B)** The longevity and P-death phenotypes of *nmur-1(ok1387)* single mutants that were rescued in *nmur-1*-expressing cells **(A)** or in sensory neurons alone **(B)**. **(C-D)** The longevity and P-death phenotypes of *daf-2(e1368)*; *nmur-1(ok1387)* double mutants, where *nmur-1* was rescued in *nmur-1*-expressing cells **(C)** or in sensory neurons alone **(D)**.

Next, we sought to determine where *nmur-1(+)* acts to influence *C. elegans* survival. Expression of *nmur-1(+)* from the sensory neuron-specific promoter *osm-6p* [21] rescued the *nmur-1* mutant survival phenotypes in wild-type and mutant *daf-2* backgrounds (Figs 4B, 4D, S4C and S4D; Tables 1, S5, S8 and S9). This result suggests that NMUR-1(+) in sensory neurons inhibits P-deaths when DAF-2 protein activity is wild type, although it has less of an effect on P-deaths when DAF-2 receptor activity is reduced (Fig 4B and 4D; Tables 1 and S9). Interestingly, NMUR-1(+) in sensory neurons rescued the non-P death phenotype of *daf-2(e1368); nmur-1(ok1387)* double mutants (Fig S4C and S4D; Tables S5 and S8) more robustly than the P-death phenotype of these animals (Fig 4D; Tables 1 and S9). Together these data suggest that wild-type NMUR-1 acts in sensory neurons to exert its multiple, context-dependent effects on *C. elegans* survival.

We also wanted to test whether LPS structure influences *nmur-1* activity to modulate P-deaths. *E. coli* CS180 has little effect on the *nmur-1* mutant phenotype in wild-type or mutant *daf-2* background (Fig 5A; Tables 1 and S10). However, a truncation of the LPS structure in *E. coli* CS2429 recapitulated the *nmur-1* mutant phenotypes on OP50: (i) an increase in P-deaths and shortening of lifespan when DAF-2 is wild type; and (ii) a decrease in P-deaths and lengthening of lifespan when DAF-2 receptor activity is reduced (Tables 1 and S10). Thus, these findings suggest that LPS structure plays a role in the NMUR-1(+)-modulation of infection-dependent P-deaths that is mediated by DAF-2 receptor protein activity.

**Fig 5.**
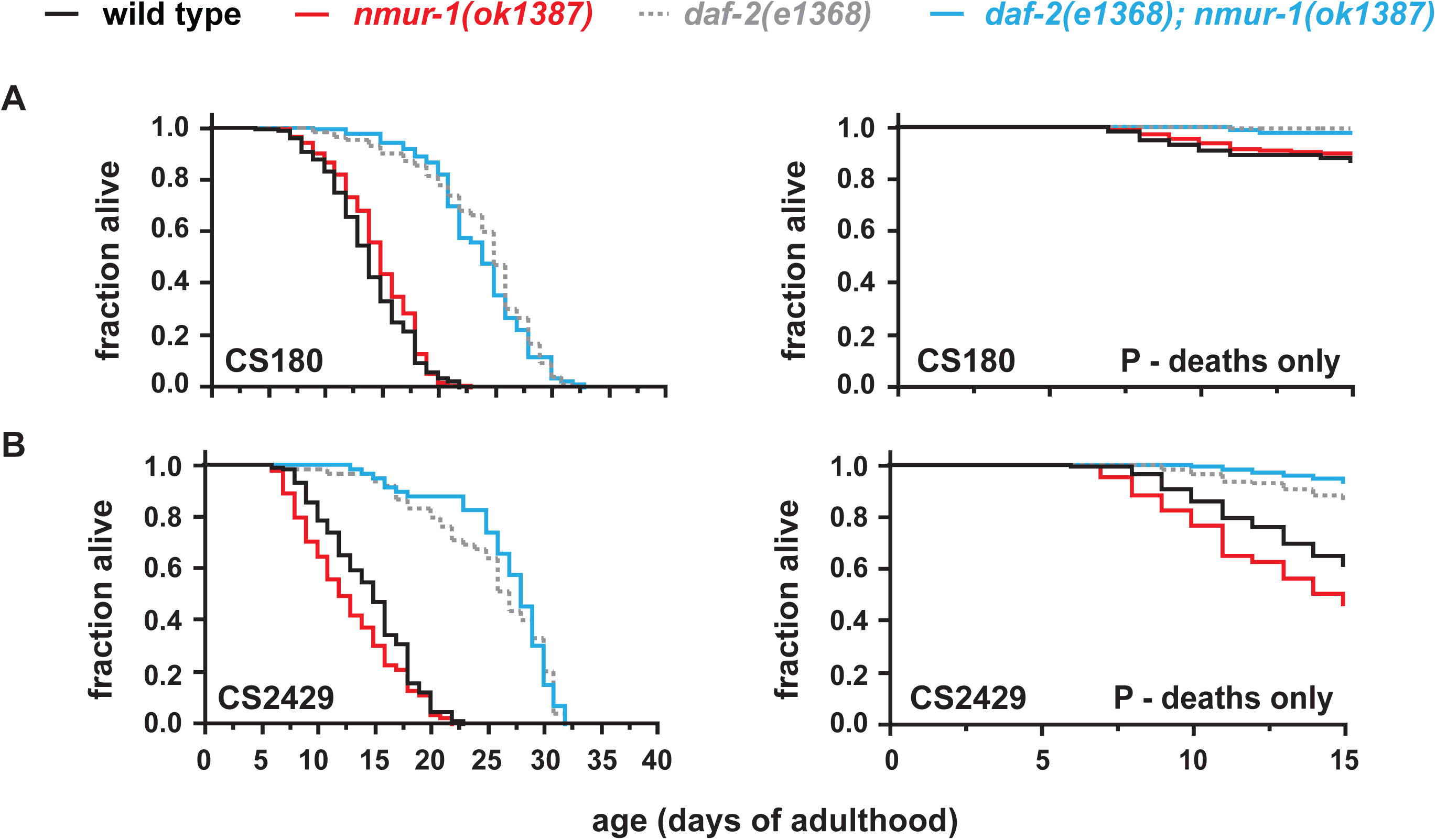
The effects of *nmur-1* depend on the bacterial LPS structure. **(A-B)** The longevity and P-death phenotypes of *nmur-1(ok1387)* single mutants and *daf-2(e1368)*; *nmur-1(ok1387)* double mutants on CS180 **(A)** versus CS2429 **(B)**.

### *nmur-1* regulates the sensory neuron expression of the insulin-like peptide *daf-28* in a context-dependent manner

The complex interactions between *nmur-1* and the *daf-2* insulin receptor motivated us to determine if they act in the same pathway. To test whether NMUR-1 acts upstream of the DAF-2 receptor, we determined if loss of *nmur-1* affects the expression of some ILPs. *C. elegans* has forty ILPs that are organized into an ILP-to-ILP network, where some ILPs have been proposed to act as agonists or antagonists of DAF-2 [28]. We focused on two ILPs, *ins-6* and *daf-28*, which encode potential DAF-2 agonists with known roles in lifespan and whose expression are modulated by bacteria-derived cues [29–32]. Because *ins-6* and *daf-28* overlap in expression [29, 32, 33] with *nmur-1* in the sensory neuron ASJ [34, 35], we compared the expression of the two ILPs in the ASJ neurons of control animals versus *nmur-1* loss-of-function mutants.

While deletion of *nmur-1* had no effect on *ins-6* expression in ASJ (Fig 6A; Table S11), it significantly increased *daf-28* expression in these neurons (Fig 6B and 6C; Table S11). This raises the possibility that *nmur-1* alters the expression of ILPs, such as *daf-28*, to exert its effects on survival. Consistent with this possibility, a deletion of *daf-28* decreased the number of P-deaths (Fig 6D; Tables 1 and S11). This suggests that NMUR-1(+) inhibits *daf-28* expression, which can subsequently suppress wild-type DAF-2 receptor function and promote survival. However, the inhibitory effect of NMUR-1(+) on *daf-28* expression only occurs when DAF-2 function is wild type (Fig 6B and 6C; Table S11), which suggests that wild-type NMUR-1 regulates the expression of certain ILPs within the context of DAF-2 receptor function. Thus, NMUR-1(+) might regulate other ILPs when the DAF-2 receptor has reduced activity.

**Fig 6.**
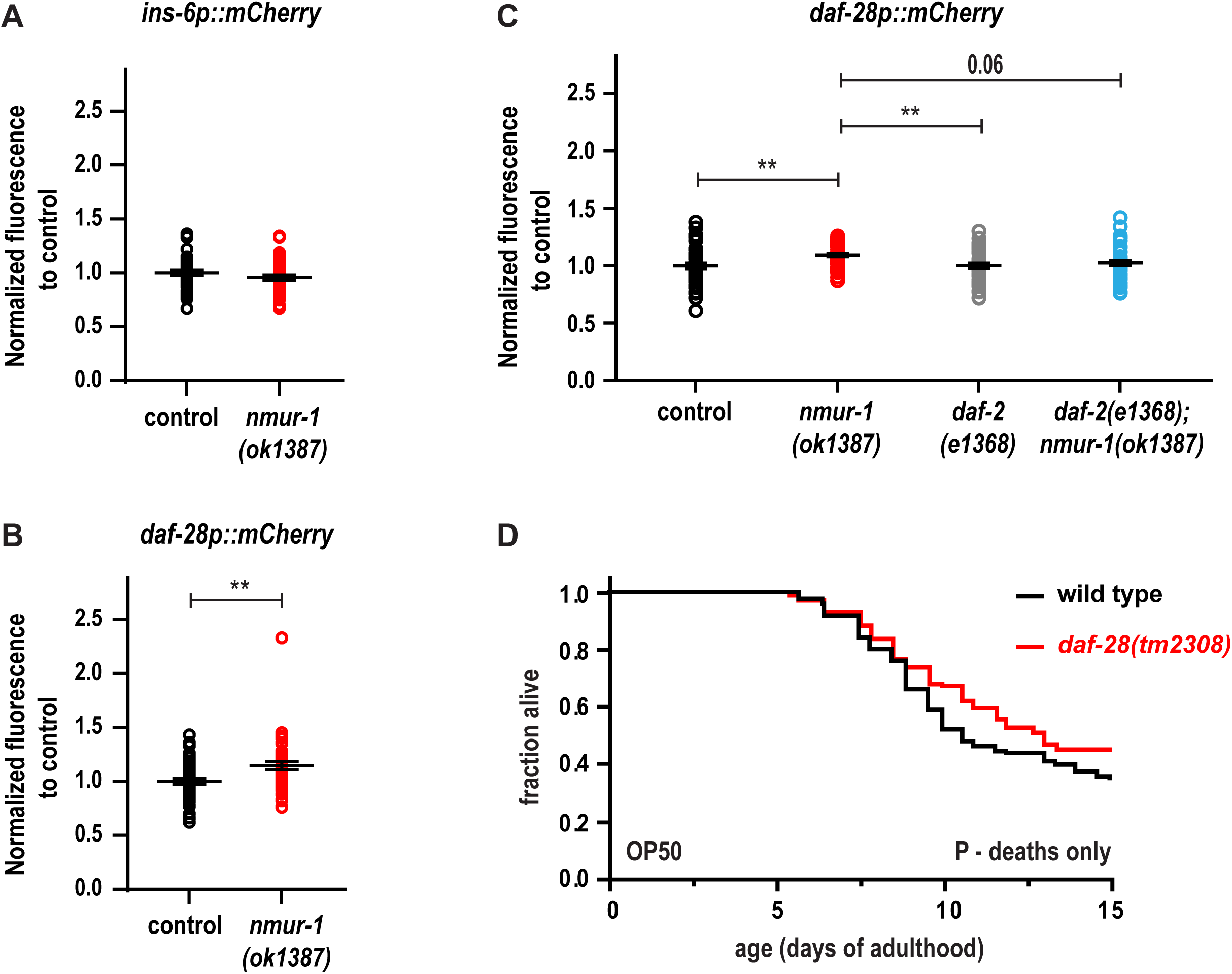
Wild-type *nmur-1* suppresses the expression of the insulin-like peptide *daf-28* in a *daf-2*-dependent manner. **(A-B)** The effects of an *nmur-1* deletion on the ASJ neuron expression of the *ins-6p::mCherry* transcriptional reporter *drcSi68* **[A**; n = 45, wild type; n = 41, *nmur-1(ok1387)*] and the *daf-28p::mCherry* transcriptional reporter *drcSi98* **[B**; n = 46, wild type; n = 44, *nmur-1(ok1387)*] at mid-L4 on OP50. **(C)** The effects of the loss of *nmur-1* on *daf-28p::mCherry* expression in ASJ neurons of mid-L4 larvae that have wild-type (black versus red circles) or reduced (grey versus blue circles) DAF-2 function [n = 49, wild type; n = 52, *nmur-1(ok1387)*; n = 49, *daf-2(e1368)*; n = 54, *daf-2(1368); nmur-1(ok1387*)] on OP50. **(D)** The OP50-dependent P-death phenotypes of animals that have wild-type *daf-28* or a deletion in the *daf-28* gene. ** indicates *P* value ≤ 0.01.

### Wild-type NMUR-1 modulates the activity of a mutant DAF-2 receptor in a *daf-16*-dependent manner

The FOXO transcription factor DAF-16 is the downstream effector of many DAF-2 functions that include longevity (reviewed by [22]), leading us to test if *nmur-1* mutant phenotypes are also *daf-16*-dependent. As with prior work [19], we found that loss of *daf-16* increased the number of all deaths, including P-deaths (Fig 7A; Tables 1 and S12), and suppressed the effect of *daf-2* mutations on all types of deaths (Fig 7B; Tables 1 and S12). Animals that lack *nmur-1* in wild-type and mutant *daf-2* backgrounds also lived as short as *daf-16* single mutants when *daf-16* was deleted in all these animals (Fig 7A and 7B; Tables 1 and S12). This raises the possibility that wild-type NMUR-1 acts with DAF-16 in the same pathway to modulate survival.

**Fig 7.**
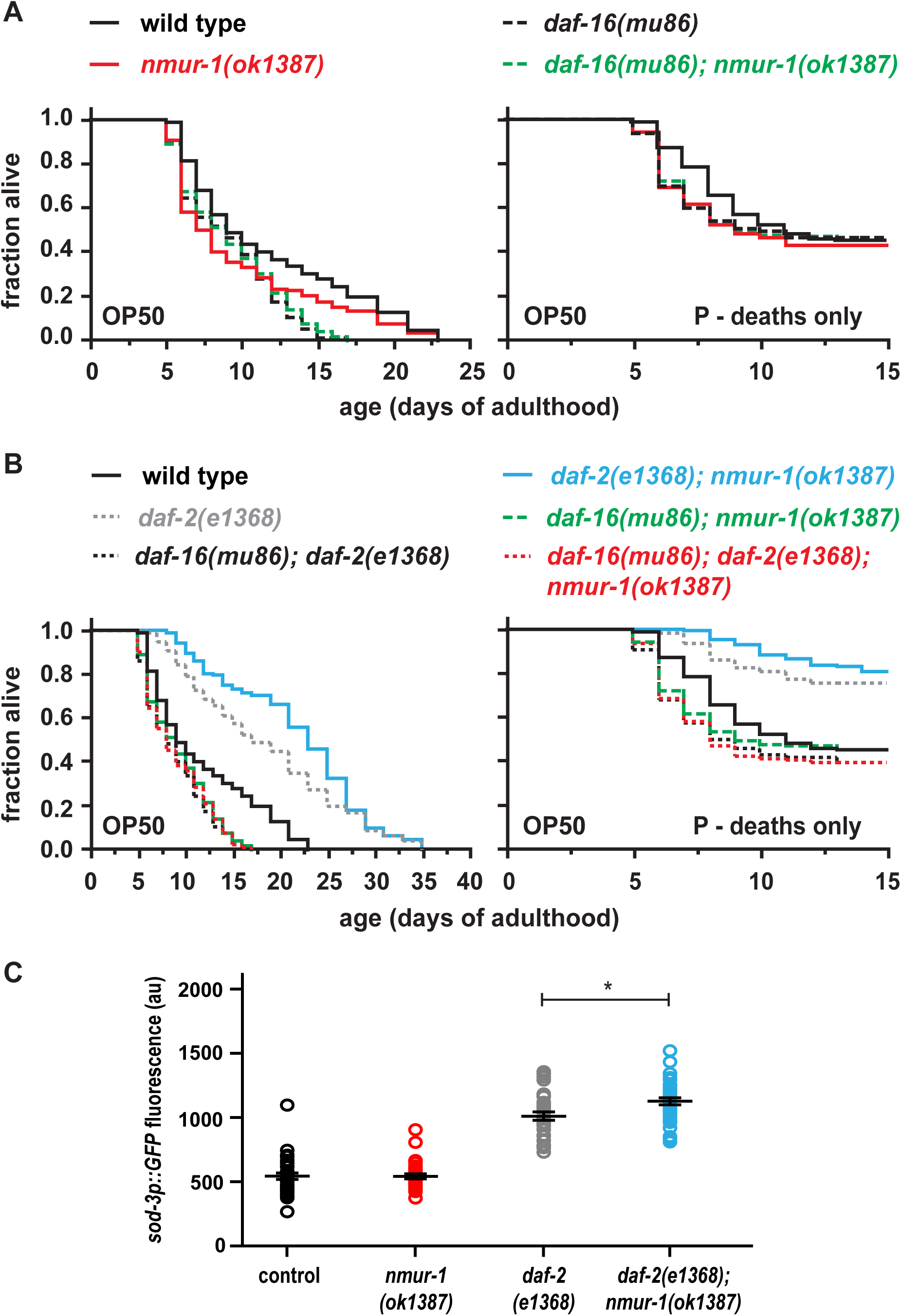
Wild-type *nmur-1* modulates the activity of a mutant DAF-2 receptor in a *daf-16*-dependent manner. **(A-B)** The *daf-16*-dependence of the longevity and P-death phenotypes of *nmur-1(ok1387)* single mutants **(A)** and of *daf-2(e1368)*; *nmur-1(ok1387)* double mutants **(B)**. **(C)** The effects of an *nmur-1* deletion on the expression of the DAF-16 target *sod-3p::GFP*, *muIs84* [36], in wild-type or mutant *daf-2* background. The quantification of *sod-3p::GFP* in the procorpus and anterior pharyngeal bulbs of one-day old animals on OP50 are shown [n = 34, wild type; n = 34, *nmur-1(ok1387)*; n = 32, *daf-2(e1368)*; n = 36, *daf-2(1368); nmur-1(ok1387*)]. * indicates *P* value ≤ 0.05, whereas “au” means arbitrary units.

To test whether NMUR-1(+) modulates DAF-16 activity, we measured the effect of the *nmur-1* mutation on the expression of a DAF-16 target gene, the manganese superoxide dismutase *sod-3*, using an integrated *sod-3p::GFP* reporter [36]. While loss of *nmur-1* alone in the presence of wild-type DAF-2 had no effect on *sod-3p::GFP*, animals that are mutant for both *daf-2* and *nmur-1* had significantly higher *sod-3p::GFP* expression than animals that are mutant for *daf-2* alone (Fig 7C; Table S12). Together these findings suggest that wild-type NMUR-1 modulates mutant DAF-2 receptor signaling by decreasing DAF-16 activity, whereas wild-type NMUR-1 might inhibit wild-type DAF-2 signaling in parallel to DAF-16 (Fig 8; see Discussion below).

**Fig. 8.**
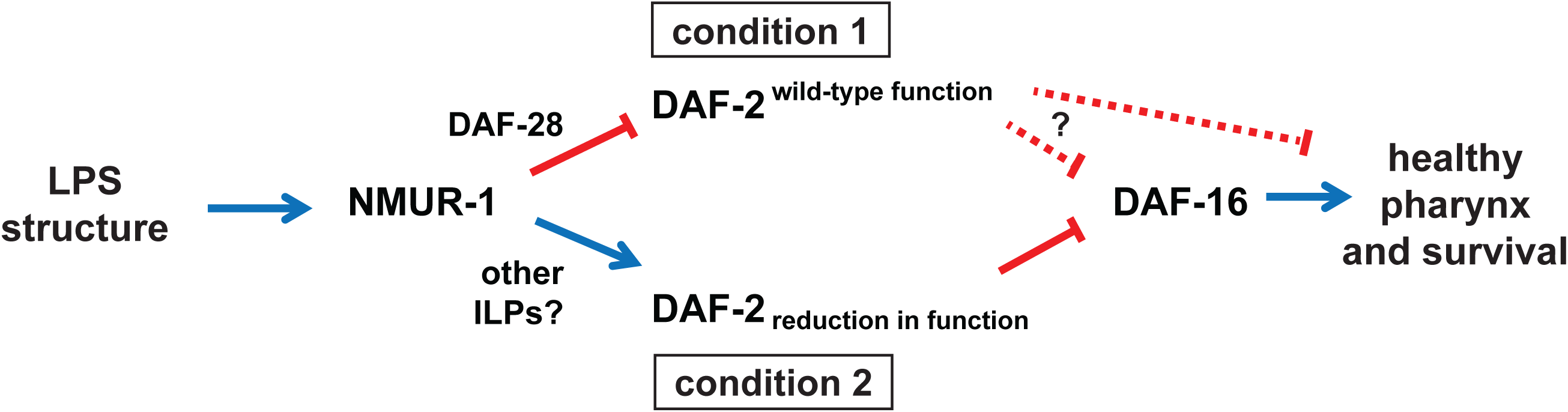
A model for how wild-type NMUR-1 adjusts DAF-2 receptor activity in regulating survival. See text for details.

## Discussion

Bacteria can modulate insulin signaling as pathogens, food, or part of the microbiome ([37]; reviewed by [38, 39]), thereby influencing key physiological processes that are important for survival. Here we used *C. elegans* genetics to dissect the contributions of these interactions on lifespan. By stratifying early and late deaths due to different bacteria-host interactions in a population, we reveal how wild-type NMUR-1 contributes to the overall survival dynamics under normal and reduced insulin signaling. Through systematic analyses of gene-gene and gene-environment interactions, our findings reveal a new role for the neuromedin U pathway in buffering the effects of perturbations to insulin signaling during bacteria-host interactions.

### Neuromedin U receptor NMUR-1 modulates survival dynamics

The survival curve of *C. elegans* is produced primarily by early deaths due to bacterial accumulation in the pharynx that are analogous to infection [15, 17] and late deaths due to other causes. Here we implicate the neuropeptide neuromedin U receptor NMUR-1 as a modulator of both early and late deaths by acting from sensory neurons. While the NMUR-1 effects on early deaths are consistent with a role in pathogen-specific innate immune responses, its effects on late deaths suggest additional role(s).

We also show that the NMUR-1 effect on early deaths occurs in response to the bacterial LPS structure. Furthermore, NMUR-1 can exert opposing effects on lifespan depending on DAF-2 receptor protein activity. More importantly, our findings on the bi-directional effects of NMUR-1 on pharyngeal-dependent survival suggest a model where NMUR-1 adjusts the dynamic range of insulin receptor signaling to promote tissue health and longevity (Fig 8).

### Bacterial LPS structure interacts with *nmur-1* to influence early death

We show that the bacterial LPS structure determines the frequency of early deaths caused by bacterial colonization of the pharynx (Figs 1E, 1G to 1J and 5; Tables 1, S1 and S10). The LPS might affect *C. elegans* pharyngeal integrity by changing pharyngeal pumping rates [7]. However, this possibility is not supported by our previous findings that wild-type animals have similar pumping rates on *E. coli* OP50, CS180 or CS2429, which have different LPS structures ([7]; and references therein). Alternatively, LPS structure might affect bacterial adherence to the pharyngeal tissues, where LPS acts as an important stimulator for the host immune system [40–43]. Some *E. coli* strains have an O-antigen that promotes adherence to tissues, an important step in pathogenesis [44]; but the strains used here lack an O-antigen (Fig 1G; [7]]. Unlike the O-antigen, the bacterial core LPS has been shown to be less adhesive, although the core LPS may regulate the expression of adherence proteins [45–47]. For example, the truncated LPS core of *E. coli* CS2198, CS2429 and OP50 might stimulate or hinder specific immune responses in *C. elegans*.

Here we find that LPS structure modulates the two activities of the *C. elegans* neuropeptide receptor NMUR-1 in altering pharynx-dependent deaths (Fig 5; Tables 1 and S10). Sun and colleagues have recently shown that *nmur-1* regulates different immune responses to specific bacterial pathogens [20]. Wild-type *nmur-1* promotes resistance to *Enterococcus faecalis*, inhibits resistance to *Salmonella enterica* and has no effect on survival on *Pseudomonas aeruginosa* [20]. While bacterial LPS has not been directly implicated in these differing responses, all three bacteria have different cell wall and LPS compositions: *E. faecalis* is a Gram-positive bacterium, which likely lacks an LPS, similar to many Gram-positive bacteria [48, 49], whereas *S. enterica* and *P. aeruginosa* are both Gram-negative bacteria with different LPS structures [50, 51]. Interestingly, the intestinal accumulation of *E. faecalis* in *nmur-1* mutants [20] is reminiscent of the bacterial colonization of pharynges on the truncated LPS mutant CS2429 (Fig. 5; Tables 1 and S10).

In rodents, LPS-induced responses are also modulated by the neuromedin U (NMU) peptide, a ligand of mammalian NMUR [52–54]. LPS exposure increases the production of the inflammatory cytokine interleukin IL-6 from peritoneal macrophages, which is abolished by the loss of the NMU peptide [53]. In this study [53], the presence of NMU promotes inflammation and LPS-induced mortality. However, in another study [54], NMU is shown to be protective against LPS-induced neuronal death, where NMU promotes the production of the neuroprotective brain-derived neurotrophic factor, BDNF, but has no effect on interleukins. While it is unclear whether the two studies used the same LPS isolate [53, 54], the NMU/NMUR signaling pathway has differing responses to LPS in both *C. elegans* and rodents. Since the NMU signaling pathway mediates LPS responses in mammals, and LPS has also been shown to affect mammalian insulin activity [55, 56], we propose that the differing NMUR-1 responses in *C. elegans* depend on the levels of insulin receptor activity (Fig 8), as we discuss below.

### *nmur-1* buffers insulin receptor signaling levels to maintain health

The insulin signaling pathway regulates *C. elegans* immune responses (reviewed by [57]), pharyngeal health, and survival [17, 19]. Severe reduction or hyperactivation of insulin receptor activity is deleterious to the animal. Insulin receptor *daf-2* null mutants exhibit lethality or embryonic and larval arrest [58], whereas a gain-of-function mutation in *daf-2* results in short-lived animals that are vulnerable to stressors [59, 60]. These studies suggest the importance of maintaining insulin receptor signaling levels at an optimal level. Modulators provide a mechanism for fine-tuning insulin receptor activity in fluctuating environments [61].

Wild-type *nmur-1*, which is co-expressed with the *daf-2* insulin receptor and/or its ILP ligands in neurons [7, 21, 34, 62], can serve as a potential modulator of DAF-2 receptor activity in regulating pharyngeal health (Fig 8). Here we show that wild-type NMUR-1 promotes healthy pharynges and prevents death (Fig 2; Tables 1 and S4), presumably by decreasing wild-type DAF-2 receptor signaling (Fig 8, condition 1). In contrast, when DAF-2 has reduced activity because of a mutation, NMUR-1 decreases the number of healthy pharynges and increases deaths (Fig 3; Tables 1 and S7), this time by potentially upregulating DAF-2 signaling (Fig 8, condition 2). These bi-directional effects of NMUR-1(+) on pharyngeal health and survival are features of neuromodulators, which ensure that cells and tissues signal within an optimal range to function appropriately across different environments [61, 63]. Here we propose that NMUR-1(+) modulates tissue and cell activities by buffering and preventing large fluctuations in DAF-2 signaling. This is supported by how NMUR-1 limits the differences in median survival between wild type and *daf-2* reduction-of-function mutants (Fig 3F; Table S7).

NMU signaling suppresses insulin secretion from *Drosophila* insulin-producing cells [23] and mammalian pancreatic β-cells in some [24–27] but not all contexts [64, 65]. In *C. elegans*, NMUR-1(+) may ensure that the insulin receptor signals appropriately by regulating the expression of specific ILP ligands. Here we show that NMUR-1(+) specifically suppresses the expression of the ILP *daf-28*, but not of *ins-6* (Fig 6A to 6C; Table S11), when DAF-2 function is wild type. Because wild-type DAF-28 increases P-deaths (Fig 6D; Tables 1 and S11), this suggests that NMUR-1(+) limits wild-type DAF-2 receptor signaling by downregulating its agonist ligand DAF-28 (Fig 8, condition 1). Intriguingly, in this context, NMUR-1(+) does not increase DAF-16 activity, as demonstrated by its lack of effect on the DAF-16 target gene *sod-3* (Fig 7C; Table S12). This suggests that NMUR-1(+) might inhibit wild-type and *daf-28*-dependent DAF-2 signaling through a mechanism parallel to DAF-16 (Fig 8, condition 1). However, DAF-16 has multiple isoforms, not all of which activate *sod-3* transcription [66]. Since the deletion mutation we used here, *mu86* (Fig 7A and 7B), removes all functional isoforms of the DAF-16 protein [66], it remains a possibility that NMUR-1(+) impedes wild-type DAF-2 signaling through a specific DAF-16 isoform.

On the other hand, when DAF-2 has a mutation-induced reduction in receptor function, NMUR-1(+) now increases its activity (Fig 8, condition 2) in a *daf-16*-dependent manner, but without altering the expression of the ILP *daf-28* (Fig 6C; Table S11). Considering that the worm has 40 ILPs with different functions [28, 29, 33, 67], it is possible that NMUR-1(+) will regulate other ILPs under this scenario.

How then does wild-type NMUR-1 determine when to promote or inhibit DAF-2 signaling? One possibility is that some DAF-2/DAF-16 targets signal back to NMUR-1 and/or its ligands. Through such a feedback mechanism, NMUR-1(+) can buffer and modulate the levels of insulin receptor signaling. Thus, in the presence of NMUR-1(+), the *C. elegans* insulin receptor is neither hyperactive nor hypoactive in response to the bacteria in the animal’s environment (Fig 8). This model highlights a mechanism that ultimately prevents large deviations in insulin pathway activity (Fig 8), which is necessary in optimizing pharyngeal health and survival.

The *C. elegans* pharynx also resembles the mammalian heart both structurally and mechanistically [68], whose health is susceptible not only to diet [69] but also to bacterial infections [70–72]. Moreover, mammalian insulin signaling plays a role in promoting cardiac health versus disease states [73–75]. Because of the high degree of conservation between *C. elegans* and mammals, we speculate that the NMUR-1-mediated buffering of insulin receptor signaling in *C. elegans* might also exist in higher animals.

## Materials and Methods

### *C. elegans* strains and growth conditions

All *C. elegans* mutants used in this study were backcrossed at least three times to wild type. Mutants that were used in the survival assays are reported in Tables 1 and S5 with their genotypes. All experiments were carried out at 25°C. However, all worms were grown for at least two generations at 20°C on the specified bacteria, before they were shifted to 25°C past the dauer larval arrest decision stage, including animals carrying the *daf-2(e1368)* or *daf-2(mu150)* mutation [58, 76, 77]. The temperature 20°C is permissive for growth for *daf-2* mutants, which prevents dauer entry, whereas 25°C is non-permissive for these animals [58].

### Bacterial strains and growth conditions

The bacterial strains that were used in the study are *E. coli* OP50, *E. coli* CS180, *E. coli* CS2198, and *E. coli* CS2429 (see [7]; and references therein). Bacterial strains were grown from single colonies in Luria-Bertani media at 37°C until the log-phase, with an optical density (OD) of ∼0.6 at 600 nm. For the experimental assays, 6-cm Nematode-Growth (NG) agar plates [16] were seeded with approximately 250 μl of bacteria and streaked to cover the entire plate (full-lawn bacterial plates). We used full-lawn plates during the lifespan assays and ILP and *sod-3p::GFP* imaging to prevent the confounding factor of worms avoiding the bacterial lawns [78]. Plates were incubated at 25°C overnight before they were used for any experiment.

### Recombining nmur-1(ok1387) away from fln-2(ot611)

To recombine *nmur-1(ok1387)* away from *fln-2(ot611)*, which is about 420 kilobases away on chromosome X of the QZ58 *C. elegans* strain, QZ58 was crossed to wild type. Among the subsequent progeny of the *nmur-1 fln-2/*+ + cross-progeny, we identified 2 recombination events out of 206 chromosomes: one progeny was homozygous for the *fln-2(ot611)* mutation and heterozygous for *nmur-1(ok1387)*; another animal was homozygous for *nmur-1(ok1387)*, but not for *fln-2(ot611)*. These animals were allowed to reproduce to isolate the *nmur-1* single mutant and the *fln-2* single mutant. The mutations were detected by PCR.

The *ok1387* deletion was detected by using the primers: ok1387 fw (5’-ATA AGT GTC ATA GAT ACA GG-3’); ok1387 rv (5’-AAT ACA TAT ACT GAT TGA CC-3’); and ok1387 int rv (5’-AAT GCT ATG GCA GAG AAG TG-3’). The mutant was detected as a 441-bp band, whereas wild type was detected as a 602-bp band.

The *ot611* point mutation was detected by using a forward primer whose 3’ end is complementary to the adenine point mutation and generates a 253-bp band with the *ot611* reverse primer, 5’-CCT GTC ACA TGA GCA CTA ATG TC-3’. The wild-type allele of *fln-2* was detected by using a forward primer whose 3’ end is complementary to cytosine and generates a 253-bp band with the *ot611* reverse primer. The presence or absence of the wild-type and *ot611* alleles were further confirmed by sequencing. We used the *ot611*_F primer, 5’-GTC ACT ATA ATA GAC GCC GTA ATG C-3’, and the *ot611* reverse primer to generate a 536-bp fragment that was sequenced to determine whether position 301 of the fragment is a C or an A.

### Lifespan assays

Worms were picked for all experiments at the late L4 stage at 25°C and were transferred onto full lawns of the specified bacteria daily for the first 6 days of adulthood, thereby preventing the mixing of subjects with their progeny. The details of the censoring during experiments are explained in the legend of Table 1. Kaplan-Meier estimates were done using the JMP 8.0.1 software (SAS). *P* values of both Wilcoxon and log-rank tests are reported in the data tables. The Wilcoxon test is the better measure of statistical significance when hazard ratios are not constant throughout an assay [7, 79], which is the case for most of our survival comparisons. For animals that carry the extrachromosomal array *ofm-1p::GFP* in Fig 6D, late L4 larvae were selected under blue light to visualize the green fluorescence.

### Necropsy analysis to determine P-deaths versus non-P deaths

The pharynges of all the dead animals in survival assays were imaged using a Nikon Eclipse Ni-U microscope and a Photometrics Coolsnap ES2 camera at 400x magnification. The surface area of the terminal pharyngeal bulb (see Fig 1C to 1E) was measured using the NIS-Elements software (Nikon Instruments, Inc). The surface area of the terminal bulb was then divided by the diameter of the body of the same animal at the region of the terminal bulb, which is also known as the grinder (area^P^/diameter^G^). This normalization addressed the possibility that the general size of the animals affected the pharyngeal surface area.

Through a principal component analysis of dead wild-type animals on OP50 (n = 387) from 8 independent survival assays, we initially separated these animals into two clusters—one with swollen pharynges (P-deaths) and one without swollen pharynges (non-P deaths). Since P-deaths happen early in the lifespan of the population [15], we used area^P^/diameter^G^ and the age of death as variables. The principal component analysis was carried out in the R 4.4.2 software [80], where we plotted the data (Fig S1) using ggplot2 [81] and ggfortify [82]. From Fig S1, we determined the threshold area^P^/diameter^G^ that would separate the two clusters, which was a ratio value of 27. This threshold value was then used to categorize animals that died with significant pharyngeal swelling (P-deaths) or with no pharyngeal swelling (non-P deaths) in all experiments. To assess the amount of P deaths only, all non-P deaths were censored in the survival assays. To assess the amount of non-P deaths, all P deaths were in turn censored.

### Imaging ILP*::mCherry* expression

*Generation of the ILP::mCherry transcriptional reporter.* The generation of the *ins-6p::mCherry* reporter *drcSi68* is as previously described [33]. The *daf-28p::mCherry drcSi98* was generated by flanking the *mCherry* gene with 3.3-kb sequences upstream of the *daf-28* start codon and 4.7-kb sequences downstream of the *daf-28* stop codon. Both 5’ and 3’ *cis* regulatory sequences of *daf-28* were amplified from YAC Y116F11 with Phusion DNA polymerase and then cloned into the pCR-Blunt vector, which was sequenced for confirmation. The subsequent reporter was next cloned into a MosSCI vector for integration (pQL184) at the ttTi4348 site of chromosome I.

*Live imaging of worms.* Animals were grown on full lawns of OP50 at 20°C, before they were shifted to 25°C at the second larval stage (L2). Worms were then imaged at 1000x magnification, once they reached the mid-L4 stage at 25°C, using a Nikon Eclipse Ni-U microscope and a Photometrics Coolsnap ES2 cooled digital camera. We quantified fluorescence intensities using a built-in fluorescence quantification algorithm (NIS-Elements, Nikon Instruments, Inc). The Student’s t-test was used to compare each ILP expression between wild type and *nmur-1(ok1387)* single mutants in Fig 6A and 6B. To compare *daf-28p::mCherry* expression in wild type, *nmur-1(ok1387)* single mutants, *daf-2(e1368)* single mutants and *daf-2(e1368); nmur-1(ok1387)* double mutants, which were imaged in parallel (Fig 6C), one-way ANOVA and Tukey’s correction were used.

### Imaging *sod-3p::GFP* expression

Animals that have an integrated *sod-3p::GFP* transgene, *muIs84* [36], were also grown on full lawns of OP50 at 20°C, before they were shifted to 25°C at the L2 stage. Worms were then imaged at 400x and 100x magnification, once they reached the first day of adulthood at 25°C, using a Nikon Eclipse Ni-U microscope and a Photometrics Coolsnap ES2 camera. Using the same NIS-Elements algorithm as above, we quantified the fluorescence intensities in the procorpus and the anterior pharyngeal bulb at 400x magnification (Fig 7C), since these tissues have the brightest fluorescence in all the animals imaged. Statistical comparisons across the multiple groups of animals were determined by one-way ANOVA, followed by Tukey’s correction.

### Statistical analyses

Statistical analyses were performed using JMP 8.0.1 (SAS) for all survival assays; GraphPad Prism 8 software for the ILP and *sod-3p::GFP* imaging measurements; and R 4.4.2 for the principal component analyses of the swollen pharynx-dependent deaths. For more details, refer to above and the figure and table legends.

## Acknowledgements

We thank the *Caenorhabditis* Genetics Center (funded by NIH P40 OD010440), I. Beets, C. Kenyon and J. Watteyne for strains used in this study. We also thank A. Caballero, D. A. Fernandes de Abreu and C. Liu for technical support in generating the *drcSi98* strain, M. Friedrich for comments on the manuscript, and E. Gourgou for supporting B.P. during the revision experiments for this manuscript. This work was supported by an ERC Starting Investigator Grant (NeuroAge 242666) and a Research Councils UK Fellowship to Q. C., and by the Novartis Research Foundation, Swiss National Science Foundation (31003A_134958), Wayne State University, the Alcedo family and NIH (R01 GM108962) to J. A.

