## Supplementary figures and images for "THE NEUROPEPTIDE NEUROMEDIN U RECEPTOR NMUR-1 BUFFERS INSULIN RECEPTOR SIGNALING IN BACTERIA-DEPENDENT *C. ELEGANS* SURVIVAL"

### Fig S1

Sifoglu et al., Figure S1

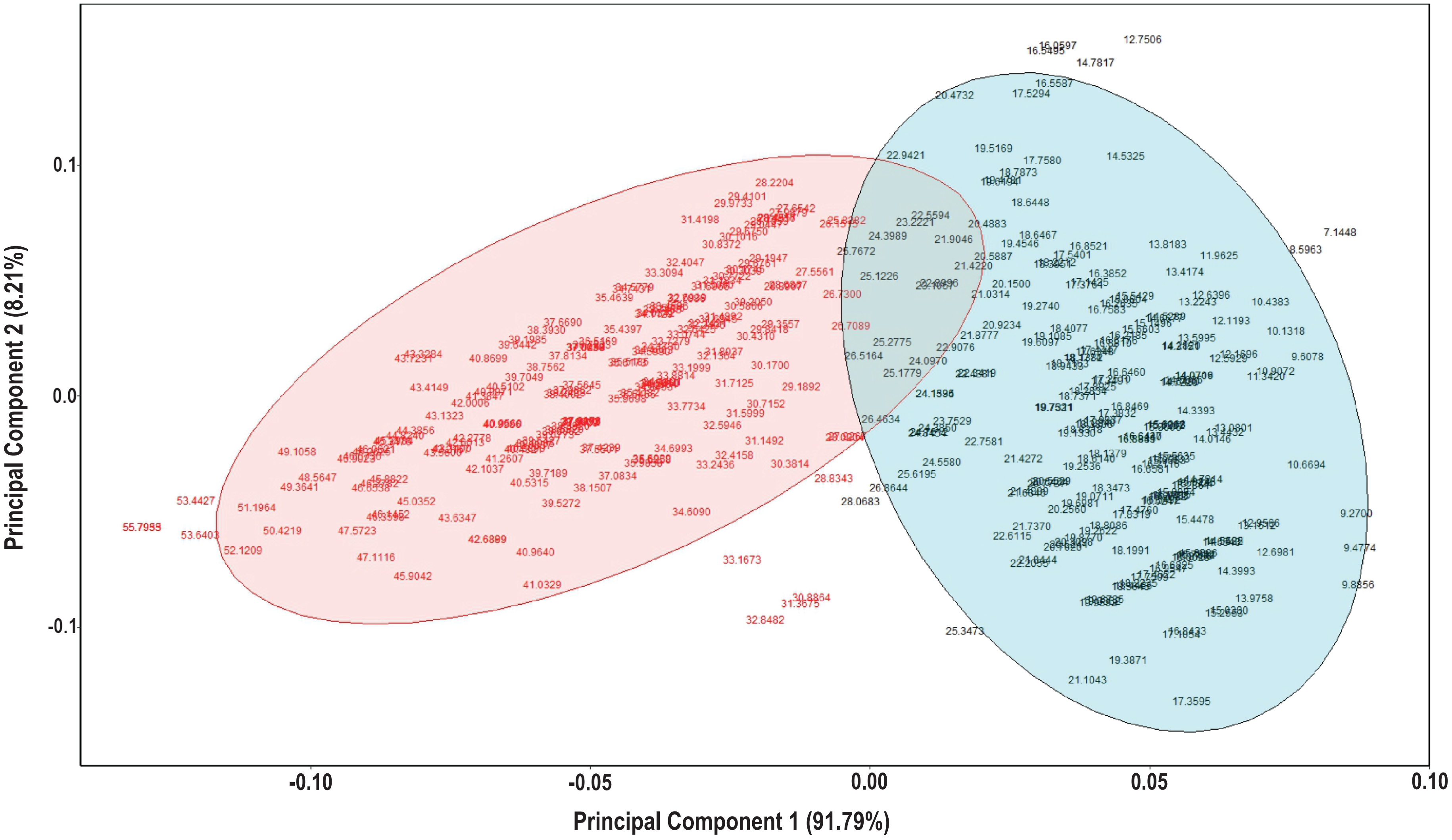

### Fig S2

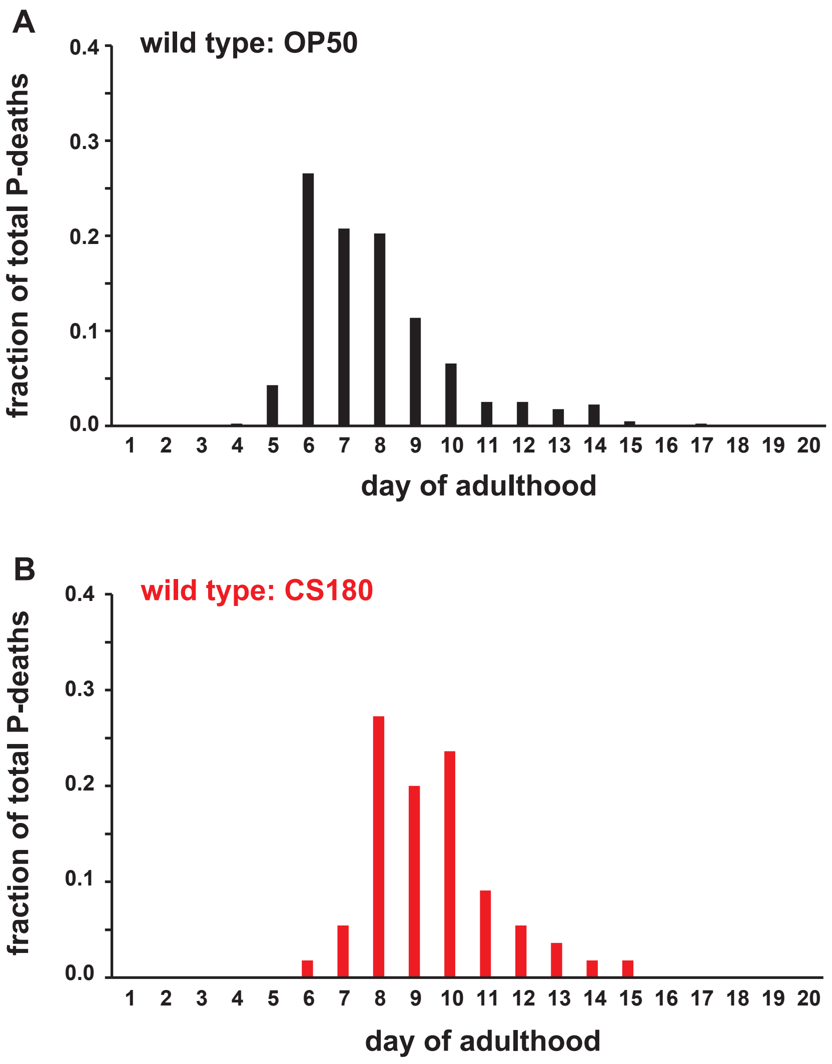

### Fig S3

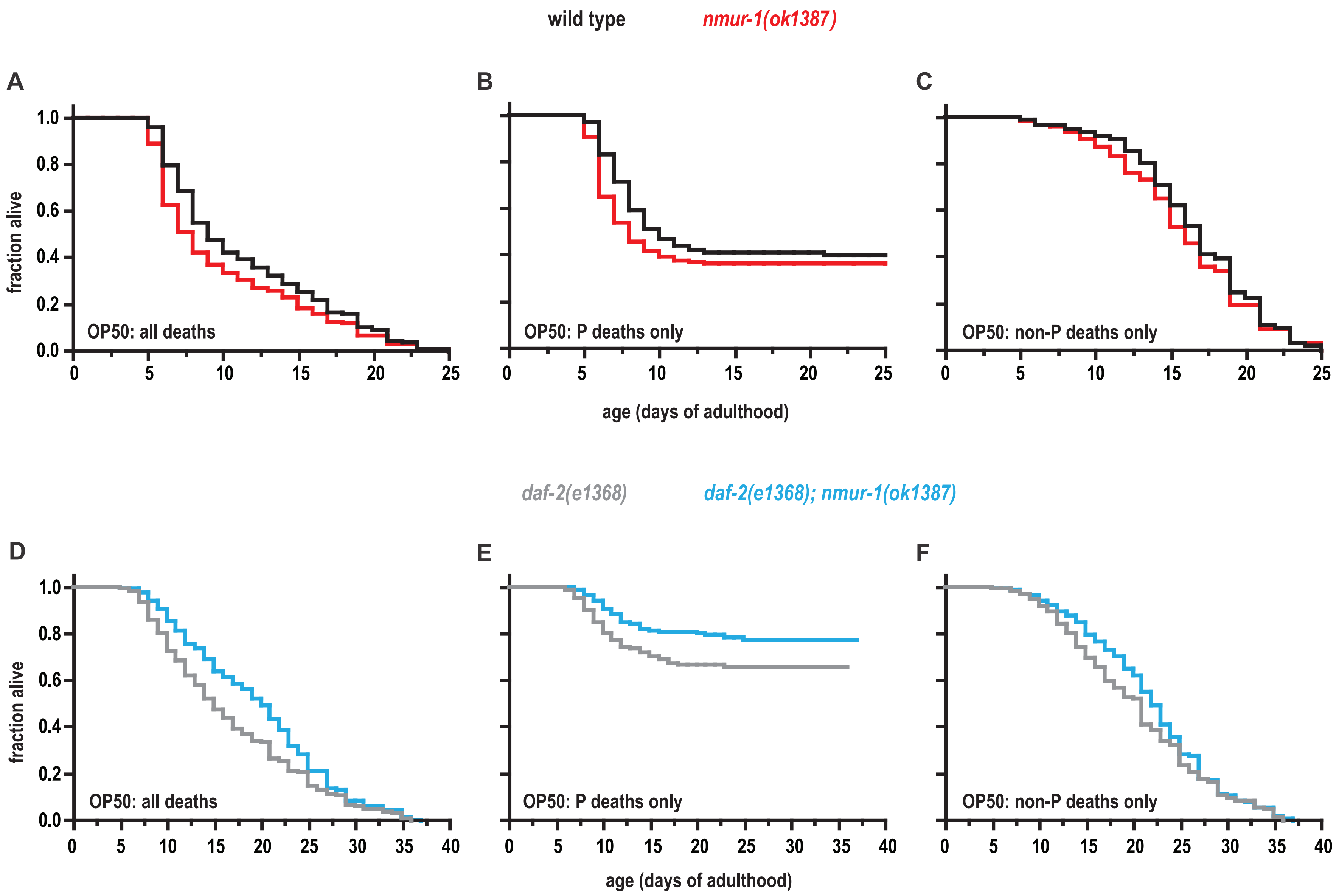

### Fig S4

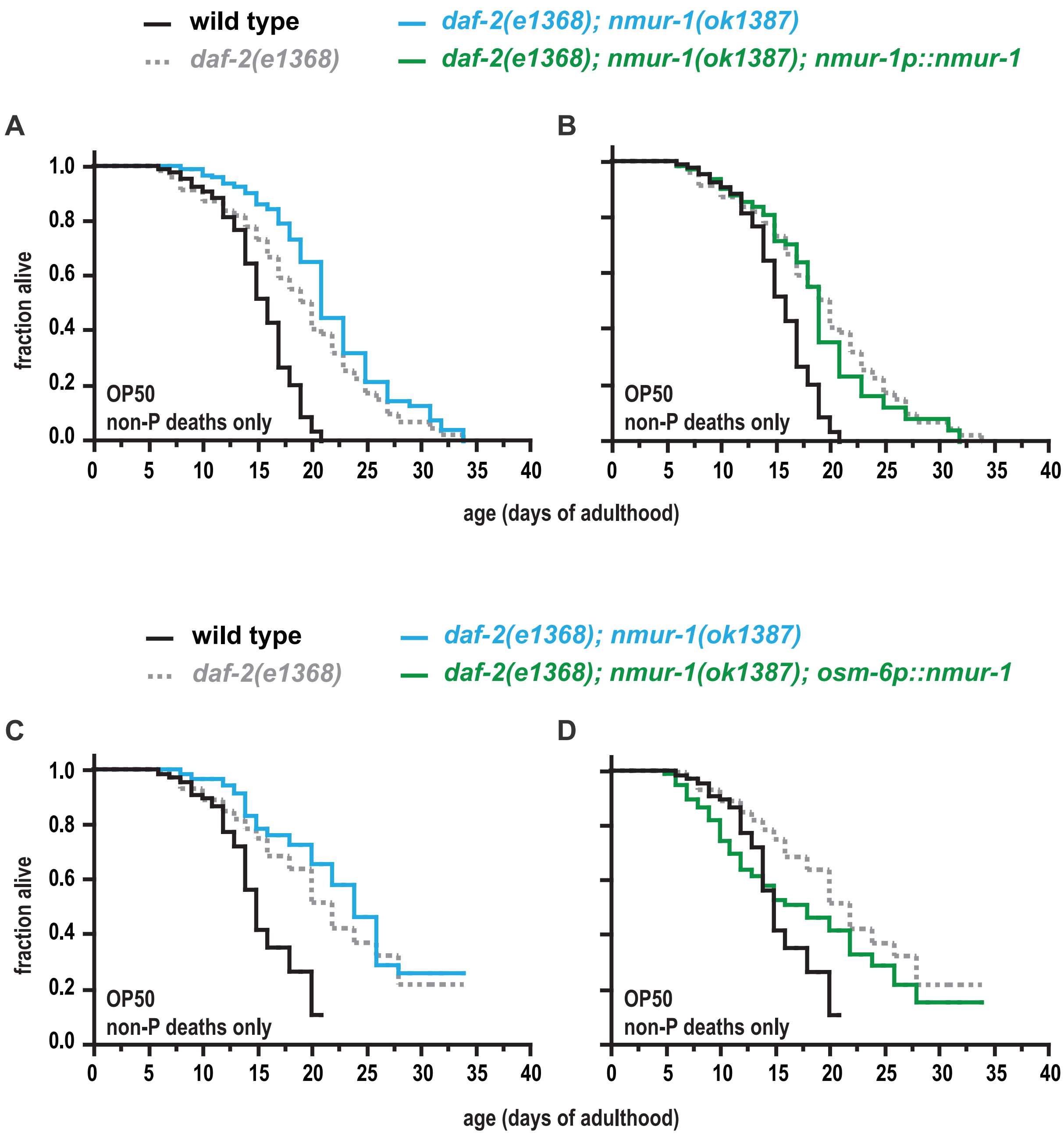
