## Supplementary material for "THE NEUROPEPTIDE NEUROMEDIN U RECEPTOR NMUR-1 BUFFERS INSULIN RECEPTOR SIGNALING IN BACTERIA-DEPENDENT *C. ELEGANS* SURVIVAL": Table S5

Table S5. *nmur-1*-dependent P-deaths versus non-P deaths on OP50

| Strain | Mean Lifespan ± SEM (Days) | # Animals Observed/ Total Initial Animals (# Trials) | *P* vs specified group (Logrank) | *P* vs specified group (Wilcoxon) | Fig |
|---|---|---|---|---|---|
| *all deaths* | | | | | |
| Wild type (WT) | 11.3 ± 0.2 | 580/1052 (7) | - | - | S3A |
| *nmur-1(ok1387)* | 10.1 ± 0.2 | 675/1052 (7) | ***< 0.0001$^{WT}$*** | ***< 0.0001$^{WT}$*** | S3A |
| *daf-2(e1368)* | 16.8 ± 0.4 | 375/1112 (6) | - | - | S3D |
| *daf-2(e1368); nmur-1(ok1387)* | 19.6 ± 0.4 | 350/892 (6) | ***< 0.0001$^{daf-2}$*** | ***< 0.0001$^{daf-2}$*** | S3D |
| *P deaths only* | | | | | |
| Wild type (WT) | | 392/1052 (7) | - | - | S3B |
| *nmur-1(ok1387)* | | 481/1052 (7) | ***< 0.0001$^{WT}$*** | ***< 0.0001$^{WT}$*** | S3B |
| *daf-2(e1368)* | | 164/1112 (6) | - | - | S3E |
| *daf-2(e1368); nmur-1(ok1387)* | | 104/892 (6) | ***< 0.0001$^{daf-2}$*** | ***< 0.0001$^{daf-2}$*** | S3E |
| *non-P deaths only* | | | | | |
| Wild type (WT) | 16.7 ± 0.3 | 188/1052 (7) | - | - | S3C |
| *nmur-1(ok1387)* | 15.9 ± 0.3 | 194/1052 (7) | 0.08$^{WT}$ | 0.09$^{WT}$ | S3C |
| *daf-2(e1368)* | 20.4 ± 0.5 | 206/1112 (6) | - | - | S3F |
| *daf-2(e1368); nmur-1(ok1387)* | 21.9 ± 0.4 | 244/892 (6) | ***0.01$^{daf-2}$*** | ***0.004$^{daf-2}$*** | S3F |
| *non-P deaths in the rescue experiments* | | | | | |
| *Rescue of daf-2(e1368); nmur-1(ok1387) double mutants with nmur-1p::nmur-1* | | | | | |
| Wild type (WT) | 15.4 ± 0.3 | 90/416 (3) | - | - | S4A,B |
| *daf-2(e1368)* | 19.3 ± 0.6 | 118/416 (3) | ***< 0.0001$^{WT}$*** | ***0.05$^{WT}$*** | S4A,B |
| *daf-2(e1368); nmur-1(ok1387)* | 21.8 ± 0.6 | 92/496 (3) | ***< 0.0001$^{WT}$***<br>***0.0004$^{daf-2}$*** | ***< 0.0001$^{WT}$***<br>***< 0.0001$^{daf-2}$*** | S4A |
| *daf-2(e1368); nmur-1(ok1387); nmur-1p::nmur-1* | 18.8 ± 0.5 | 112/496 (3) | ***< 0.0001$^{WT}$***<br>0.6$^{daf-2}$<br>***< 0.0001$^{ok1387*}$*** | ***0.007$^{WT}$***<br>0.8$^{daf-2}$<br>***< 0.0001$^{ok1387*}$*** | S4B |
| *Rescue of daf-2(e1368); nmur-1(ok1387) double mutants with osm-6p::nmur-1* | | | | | |
| Wild type (WT) | 15.3 ± 0.4** | 66/336 (2) | - | - | S4C,D |
| *daf-2(e1368)* | 20.7 ± 0.6** | 71/336 (2) | ***< 0.0001$^{WT}$*** | ***0.003$^{WT}$*** | S4C,D |
| *daf-2(e1368); nmur-1(ok1387)* | 22.3 ± 0.5** | 73/336 (2) | ***< 0.0001$^{WT}$***<br>***0.05$^{daf-2}$*** | ***< 0.0001$^{WT}$***<br>***0.006$^{daf-2}$*** | S4C |
| *daf-2(e1368); nmur-1(ok1387); osm-6p::nmur-1* | 17.8 ± 0.7** | 88/336 (2) | 0.5$^{WT}$<br>***0.002$^{daf-2}$***<br>***< 0.0001$^{ok1387*}$*** | ***0.04$^{WT}$***<br>***< 0.0001$^{daf-2}$***<br>***< 0.0001$^{ok1387*}$*** | S4D |
